# T cells in testicular germ cell tumors: new evidence of fundamental contributions by rare subsets

**DOI:** 10.1101/2023.09.28.559963

**Authors:** Rashidul Islam, Jannis Heyer, Miriam Figura, Xiaoyan Wang, Xichen Nie, Benedict Nathaniel, Sivanjah Indumathy, Katja Hartmann, Christiane Pleuger, Monika Fijak, Sabine Kliesch, Florian Dittmar, Adrian Pilatz, Florian Wagenlehner, Mark Hedger, Bruce Loveland, James H. Hotaling, Jingtao Guo, Kate Loveland, Hans-Christian Schuppe, Daniela Fietz

**Affiliations:** Dept. of Veterinary Anatomy, Histology and Embryology, Justus Liebig University, Giessen, Germany; Centre for Reproductive Health, Hudson Institute for Medical Research, Clayton, Victoria, Australia; Department of Molecular and Translational Sciences, Monash University, Clayton, VIC, Australia; Dept. of Urology, Pediatric Urology and Andrology, Justus Liebig University, Giessen, Germany; State Key Laboratory of Reproductive Biology, Institute of Zoology, Chinese Academy of Sciences, Beijing, China; Beijing Institute of Stem Cell and Regenerative Medicine, Beijing, China; Division of Urology, Department of Surgery, University of Utah School of Medicine, Salt Lake City, Utah, USA; Hessian Centre of Reproductive Medicine, Justus-Liebig-University, Giessen, Germany; Institute of Anatomy and Cell Biology, Justus Liebig University, Giessen, Germany; Centre of Reproductive Medicine and Andrology, University of Muenster, Muenster, Germany; Burnet Institute, Melbourne, Australia

## Abstract

**BACKGROUND:** Immune cell infiltration is heterogeneous but common in testicular germ cell tumors (TGCT) and pre-invasive germ cell neoplasia in situ (GCNIS). Tumor-infiltrating T cells including regulatory T (Treg) and follicular helper T (Tfh) cells are found in other cancer entities, but their contributions to TGCT are unknown.

**METHODS:** Human testis specimens from independent patient cohorts were analyzed using immunohistochemistry, flow cytometry and single-cell RNA sequencing (scRNA-seq) with special emphasis on delineating T cell subtypes.

**RESULTS:** Profound changes in immune cell composition within TGCT, shifting from macrophages in normal testes to T cells plus B and dendritic cells in TGCT, were documented. In most samples (96%), the CD4+ T cell frequency exceeded that of CD8+ cells, with decreasing numbers from central to peripheral tumor areas, and to tumor-free, contralateral testes. T cells including Treg and Tfh were most abundant in seminoma compared to mixed tumors and embryonal carcinoma.

**CONCLUSION:** Despite considerable heterogeneity between patients, T cell subtypes form a key part of the TGCT microenvironment. The novel finding of rare Treg and Tfh cells in human testis suggests their involvement in TGCT pathobiology, with implications for understanding tumor progression, to assess patients’ prognosis, and as putative targets for personalized immunotherapy.

## BACKGROUND

Testicular cancer is the most common solid tumor in young men between the age of 15-45 years and accounts for approximately 1% of newly diagnosed male cancers worldwide (1, 2). The overall incidence of testicular cancer is steadily increasing globally, but the underlying causes are not well understood (3). In 90-95% of cases, testicular neoplasms arise from gonocytes that are understood to first form pre-invasive germ cell neoplasia in situ (GCNIS) cells, then progress to form testicular germ cell tumor (TGCT) classified as either seminomas or non-seminomas (1, 2). Seminomas account for ∼60% of total TGCT and are characterized as homogeneous tumors with the appearance of atypical gonocytes that are blocked at the earliest stage of differentiation, while the heterogeneous non-seminomatous tumors can manifest as embryonal carcinoma (the most undifferentiated type), yolk sac tumor, choriocarcinoma or teratoma (1, 4, 5). It remains largely unclear under which circumstances GCNIS develops into seminoma or non-seminoma, seminoma transitions into non-seminoma, and which factors drive tumor progression including metastatic spread (1, 2, 6). The infiltration of immune cells has been identified as a hallmark feature of TGCT (7-9). Hence, profound changes in the landscape of testicular immune cells, and their interactions with germ cells and other testicular somatic cells, likely contribute to a unique tumor microenvironment (TME) in TGCT (10, 11).

Under physiological conditions, immune cells are essential for maintaining “immune privilege” within the testis. Macrophages are normally the most abundant immune cells in the adult testis, displaying an anti-inflammatory phenotype, including anergy to inflammatory stimuli, with a phenotype shaped by the local microenvironment (12, 13). Present in small numbers, antigen-presenting dendritic cells (DC) and regulatory T lymphocytes (Treg) are important for the induction of systemic tolerance to developing germ cell antigens (13, 14). The physiological milieu is also determined by a combination of immunoregulatory and immunosuppressive factors, including cytokines such as TGF-β and interleukin (IL)-10, activin A, and androgens, provided by resident immune cells and somatic Sertoli, Leydig and peritubular cells (11, 15). This delicate milieu can be disrupted by pathogens such as bacteria and viruses, as well as by the presence of neoplastic germ cells (10, 13).

T cells represent the majority of tumor infiltrating lymphocytes (TIL) in TGCT, especially in seminomas, with increased numbers of macrophages and DC (7-9, 16-19). TIL can include abundant B cells; these may form follicular-like structures (FLS) that resemble germinal centres found in tertiary lymphoid organs (8, 19, 20). Despite histopathological similarities in early cancer stages, the composition of immune cell infiltrates in TGCT markedly differs from low-grade testicular inflammation in infertile men (8). Previous studies report significantly increased levels of transcripts encoding pro-inflammatory and T helper cell type (Th) 1-driven cytokines and B cell supporting chemokines in GCNIS and seminoma, compared to testes with normal phenotype or hypospermatogenesis (HYP) with focal inflammatory lesions (8, 20, 21). Interestingly, *IL-6*, encoding the pro-inflammatory cytokine IL-6, was not only found to be upregulated in TGCT, but its expression level was directly associated with the metastatic status of seminomas (22). Apart from promoting antibody production by B cells (23) and polarization of alternatively activated macrophages (24), IL-6 also regulates the differentiation of CD4+ Th and CD8+ cytotoxic T (Tc) cells into their functionally diverse subtypes (11, 25-27). In other malignant tumors such as breast, ovarian, colorectal, or lung cancers, TIL have been associated with versatile effects on tumor growth, metastatic behavior and patient survival (28). It was shown that Treg (mainly producing immunosuppressive cytokines such as TGF-β, IL-10, and IL-35) infiltrate the TME and are critically related to poor prognosis in various cancers (29-31). Follicular helper T (Tfh) cells located in lymphoid organs, support B cells and drive immunoglobulin class-switching and affinity maturation in germinal centres (32). For a range of cancers, the presence of Tfh was generally positively associated with long-term survival (33, 34), except for hepatocellular carcinoma (34, 35).

A recent transcriptome analysis using different regions of TGCT revealed a marked heterogeneity of differentially expressed genes in tumor-central versus tumor-adjacent regions with most of the differentially expressed genes falling into the category of immune-related processes, indicating the importance of immune cells in TGCT progression and metastatic spread (4). Immune profiling in primary and metastatic TGCT identified activated T cell infiltration, increased programmed cell death-1/ligand-1 (PD-1/PD-L1) spatial interaction, and low percentages of Treg cells in seminomas associated with a good prognosis, while high neutrophil and macrophage signatures were observed in non-seminomas. Advanced TGCT stage was associated with decreased pan-T cell and NK cell signatures, while Treg, neutrophil, mast cell, and macrophage signatures were increased (9). However, clinical trials of immune checkpoint inhibitors for TGCT treatment have failed, reflecting an incomplete understanding of the testis-specific immune microenvironment (36).

This study aimed to delineate the variations and consistencies of immune cell profiles in TGCT. We comprehensively analyzed the landscape of infiltrating immune cells in testis samples from distinct patient cohorts using immunohistochemistry and flow cytometry, underpinned by transcriptome analyses using single-cell RNA sequencing (scRNAseq) datasets.

## MATERIALS AND METHODS

### Patients

Characterization of immune cell populations in the human testis was first performed on archived tissue specimens obtained from men who underwent surgery for suspicion of testis cancer or multifocal biopsy for testicular sperm extraction indicated by obstructive or non-obstructive azoospermia This retrospective patient cohort (n=81, 20-69 years, median age 36 years) originated from the Department of Urology, Pediatric Urology and Andrology at Giessen University Hospital and the Department of Clinical Andrology, Center for Reproductive Medicine and Andrology at the University of Münster, Germany.

As part of a prospective TGCT cohort study established at the Department of Urology, Pediatric Urology and Andrology at Giessen University Hospital, testicular tissue including fresh samples was obtained from different locations of tumor-bearing as well as from contralateral testes (n=24 patients, 23-56 years, median age 35 years). In addition to this study population, testicular tumor biopsies were provided by the University of Utah for scRNA-seq (n=4 patients).

All patients gave their written informed consent to the use of tissue samples for research purposes. This procedure was approved by the ethics committee of the Medical Faculty of the Justus Liebig University Giessen (Ref. No. 26/11; 152/16) and the University of Utah (IRB approved protocol #00075836).

### Histological evaluation

For classification of human testicular tissues, specimens were fixed by overnight immersion in Bouin’s solution and embedded in paraffin. Sections (5 μm) were stained with hematoxylin and eosin (H&E). The histopathological assessment included a comprehensive assessment of immune cell infiltrates (8); it was combined with a score count analysis of spermatogenesis to classify non-cancerous testicular specimens (37). Based on the pathologcal findings reported during uro-oncological work-up, patients with TGCT were categorized into three groups: (I) pure seminoma, (II) embryonal carcinoma (≥80% of tumor tissue), and (III) mixed tumors. A histopathological analysis of the TGCT tissue sections from different locations in tumor-bearing and contralateral testes was performed as reported earlier (8).

### Immunohistochemical analysis

For IHC, five sample categories from the retrospective patient cohort were used: normal spermatogenesis (NSP) (n=10), HYP with lymphocytic infiltrates (HYP+ly; n=11), GCNIS (n=14), GCNIS with lymphocytic infiltrates (GCNIS+ly; n=12), seminoma (n=24), and embryonal carcinoma (n=10). IHC was performed as previously described (8). Briefly, 5μm tissue sections were deparaffinized and rehydrated. Heat-mediated antigen retrieval was performed by immersing slides in Tris-EDTA buffer (10 mM Tris Base, 1 mM EDTA, 0.05% Triton X-100, pH 9.0) for 20 min at 522 W in a microwave oven. To inhibit endogenous peroxidase activity, the slides were incubated in 3% hydrogen peroxide in TBST (Tris-buffered saline + 0.1% Triton X-100, pH 7.6) for 15 min at room temperature (RT), followed by washes with TBST. Sections were next incubated in 1.5% bovine serum albumin (BSA) (Roth, Karlsruhe, Germany) in TBST at RT for 30 min to block non-specific binding. Sections were incubated with primary antibodies (Supplemental Table 1) overnight at 4°C in a humidified chamber. The next day, slides were washed three times (5 min each), between incubations at RT using TBST. Appropriate biotinylated secondary antibodies diluted in TBST (Supplemental Table 1) were added for 1 hour at RT. Next, the sections were incubated with VECTASTAIN Elite ABC kit, Peroxidase (standard) according to the manufacturer’s instructions (Vector Laboratories, Newark CA, USA) for 45 min at RT. Finally, immunostaining was visualized using ImmPACT AEC Substrate Kit, Peroxidase (Vector Laboratories) for < 25 min or Vector NovaRED Peroxidase Substrate (Vector Laboratories) for < 10 min. Sections were lightly counterstained with Mayer’s hematoxylin (Roth) for detection of cytoplasmic proteins. No counterstain was applied for detection of nuclear proteins to avoid signal masking. Sections stained with NovaRED were dehydrated and mounted with Kaiser’s glycerol gelatine (phenol-free; Roth) or Eukitt® quick-hardening mounting medium (Sigma-Aldrich, St. Louis MO, USA). Conditions used for all antibodies were optimized using human tonsil tissue as the positive control. IHC image analysis was performed at Monash University, Australia (Aperio ScanScope AT Turbo at Monash Histology Platform facility). Images were analyzed using the Aperio ImageScope (V12.4.3.5008) software.

### Flow cytometric analysis of immune cells

Fresh tissue samples were collected from different areas of the tumor-bearing testis (tumor center [“Tumor”], adjacent to tumor [“Tumor-Adj”], distant from tumor [“Tumor-Dis”]) and contralateral testes ([“Contralateral 1”] and [“Contralateral 2”]) (Supplemental Fig. 1A). Tissue pieces were immediately transferred into RPMI medium on ice for flow cytometry, and adjacent tissues from the same location were collected and fixed separately for histological evaluation and IHC analysis. Single cell suspensions for flow cytometric analysis were generated using a previously established protocol (38). First, the samples were minced into small pieces and then enzymatically digested with collagenase D (1.5 mg/ml, Sigma-Aldrich) for 45 min at 37° C in a shaking heater at 800 rpm. Afterwards, single cells were retrieved by filtration using a 100 μM cell strainer (Greiner Bio-One GmbH, Kremsmünster, Austria). Single cell pellets were re-suspended with PBS, incubated with Viobility™ 405/520 Fixable Dye (Miltenyi Biotec, Bergisch Gladbach, Germany) according to the manufacturer’s protocol and washed with MACs Quant buffer (2mM EDTA+0.05% BSA in PBS). Cells were incubated with the antibodies directed to cell surface protein epitopes (Supplemental Table 2) according to the manufacturer’s protocol, with appropriate isotype controls for each. After washing in MACs Quant buffer, cells were incubated with FOXP3 staining buffer and then washed with permeabilization buffer (both from Miltenyi Biotec). The cell pellets were resuspended with permeabilization buffer and incubated with 20 μl FcR blocking reagent (Miltenyi Biotec) for 10 min at 4°C in the dark. Subsequently, cells were incubated with antibodies targeting intracellular antigens (Supplemental Table 2). Finally, cells were washed with 1X permeabilization buffer and re-suspended in 100-200 μl MACs Quant buffer then run in the MACSQuant® Analyzer 10 Flow Cytometer. Data were analyzed with FlowJo™ v10.8.1 and graphs generated with GraphPad Prism 9.3.1 software.

### scRNA-seq: Sample preparation, library construction and sequencing

Testis specimens (n=4) were placed into cold PBS and transported on ice. Tissues were digested following the standard two step enzymatic isolation protocol as described in (39). Briefly, specimens were digested with collagenase type IV (Sigma Aldrich) for 5 min at 37 °C with gentle agitation (250 rpm), then shaken vigorously and incubated for another 3 min. The tubules were sedimented by centrifugation at 200× g for 5 min and washed with Hanks’ Balanced Salt Solution (HBSS) before digestion with 4.5 mL 0.25% trypsin/ethylenediaminetetraacetic acid (EDTA; Invitrogen) and 4 Kunitz unit (kU) DNase I (Sigma-Aldrich). The suspension was triturated vigorously three to five times then incubated at 37 °C for 5 min. The process was repeated in 5 min increments for up to 15 min total. The digestion was stopped by adding FBS (fetal bovine serum) to a final concentration of 10% (Gibco, Billings MT, USA). Single testicular cells were obtained by filtering through 70 μm and 40 μm strainers (Thermo Fisher Scientific, Waltham MA, USA) to obtain single cell suspensions. The cells were pelleted by centrifugation at 600× g for 15 min and washed twice with PBS (Thermo Fisher Scientific). Cell number was measured using a hemocytometer, and cells re-suspended in 1× PBS/0.4% BSA (Thermo Fisher Scientific). Single cell suspensions were loaded and run on the 10x Chromium Controller using Chromium Single Cell 3′ v3.1 reagents (10x Genomics PN-1000121). Sequencing libraries were prepared following the manufacturer’s instructions, using 13 cycles of cDNA amplification, followed by an input of ∼100 ng of cDNA for library amplification using 12 cycles. The resulting libraries were then sequenced on a 2 X 100 cycle paired-end run on Illumina HiSeq 2500 or Novaseq 6000 instruments.

### scRNA-seq data analysis

10X data matrixes of normal testis (n=3; “donors”, pooled data, GEO: GSE120508) (39) and TGCT (n=4) were imported into the Seurat V4.0 R package (https://satijalab.org/seurat/) to perform analytical quality control, data normalization, dimensional reduction, data visualization etc. First, Seurat object was created for each dataset and then merged them into a single object. The following criteria were next applied to the merged datasets containing four tumor and three normal testes samples: nCount_RNA (total number of molecules detected within a cell) >1,000, nFeature_RNA (number of genes detected in each cell) between 200 and 8,000, percent.mt (mitochondrial gene percentage) < 5. After filtering, a total number of 10,153 cells remained for analysis (Supplemental Table 3). Data integration was performed to remove batch effects across different samples. The filtered matrix was normalized in Seurat v.4 with default parameters and the top 2,000 variable genes were then identified using the “vst” method in Seurat FindVariableFeatures function. Variable “nCount_RNA “and “percent.mt” were regressed out in the scaling step and Principal-Component-Analysis (PCA) was performed using the top 2,000 variable genes. Then t-distributed stochastic neighbor embedding (tSNE) was performed on the top 30 principal components for visualizing the cells. Clustering was performed on the PCA-reduced data with resolution 3 to refine the result and the clusters were identified/annotated based on expression of well-established cell-specific marker genes (39-45) throughout the 46 primary clusters (Supplemental Fig. 2 A-B). Primary clusters containing the same cell type were merged for better visualization. Secondary clustering of T cells was performed with resolution 2 to obtain the clearest outcome and then each of the 17 secondary clusters was identified based on expression of well-established T cell subtypes-specific marker genes (40) (Supplemental Fig. 3 A-B). Cell types identified as the same in the secondary clusters of T cells were merged for better visualization.

### Statistical analysis

An ordinary one-way ANOVA including Tukey’s Honest Significant Difference Test was performed for statistical analysis of different cells of the multiple comparisons across the different localization of TGCT samples by flow cytometry (Supplemental Table 4) using GraphPad Prism 9.3.1 software.

## RESULTS

### The immune cell landscape in TGCT is profoundly changed compared to controls with complete spermatogenesis

The phenotypic analysis of different immune cells in archival human testis specimens by IHC revealed marked infiltration of immune cells only in pathological (non-neoplastic and neoplastic) conditions. Representative results of IHC for each patient subgroup are shown in Fig. 1.

**Fig. 1:**
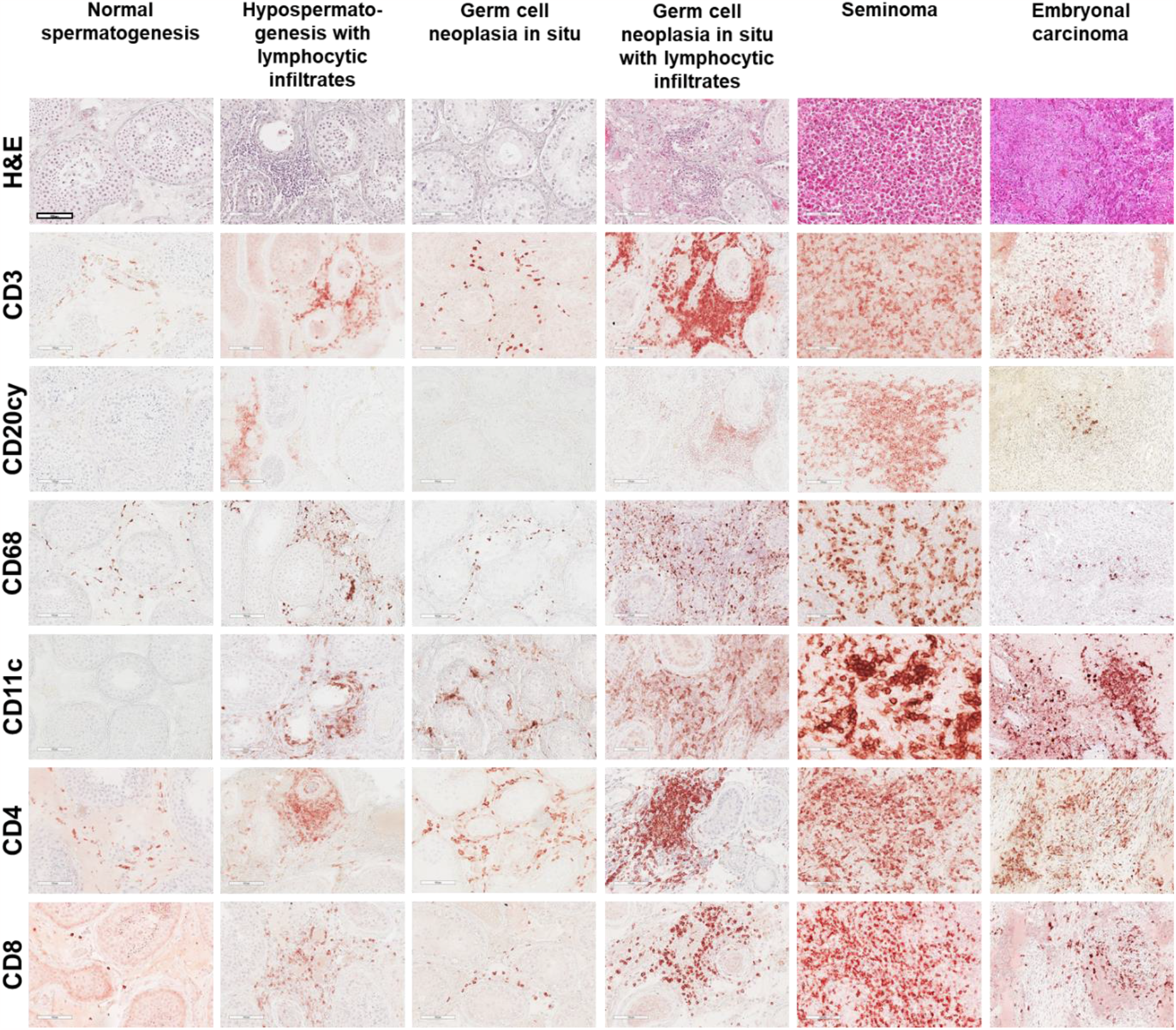
Immune cells in normal and pathological human testis. H&E staining and IHC analysis (dark red stain indicates positive signal) illustrates the prevalence and distribution of distinct immune cell populations in each pathology and can be compared to the physiological situation (NSP). The photomicrographs shown are representative for each subgroup. In general, areas of immune cell abundance were identified in TGCT (both seminoma, n=24, and embryonal carcinoma, n=10) compared to normal spermatogenesis (n=10), hypospermatogenesis with lymphocytic infiltrates (n=11), germ cell neoplasia in situ (GCNIS; n=14), GCNIS with lymphocytic infiltrates (n=12); CD3+ T cells represent the majority of immune cells in TGCT samples, especially in seminoma. White bar in top left-hand panel indicates 100μm, all images at the same magnification.

In samples with NSP, which served as the control, CD68+ macrophages were the most abundant resident immune cell, appearing distributed throughout the human testicular interstitium. CD3+ T cells, CD4+ Th and CD8+ Tc cells were detected in low numbers, particularly when compared to pathological conditions. Also, CD11c+ DC were extremely rare, and no CD20cy+ B cells were detected in NSP samples. Cells with Treg-specific markers CD25+ and FOXP3+ were scarce, and cells with Tfh-specific markers CXCR5+ and BCL6+ were absent in NSP (Supplemental Fig. 1F).

In samples with non-neoplastic low-grade testicular inflammation, seen in patients with different forms of azoospermia, infiltrating immune cells were mainly found focally or multifocally in a peritubular and/or with perivascular distribution. The most abundant cell types were CD3+ T cells, CD4+ Th, and CD8+ Tc cells, followed by CD68+ macrophages. CD20cy+ B cells and CD11c+ DCs were only rarely observed. This also applied to CD25+ and FOXP3+ cells, considered as Treg, and CXCR5+, with the latter most likely Tfh cells. No BCL6+ cells were found (Supplemental Fig. 1F).

In GCNIS and GCNIS+ly, the infiltration density greatly varied between samples. Compared to NSP, higher numbers of CD3+ T cells, CD4+ Th, and CD8+ Tc cells were found in both GCNIS subgroups, thus these represented the most prominent immune cell types with the highest density in GCNIS+ly samples. It was noted that CD4+ Th cells were relatively more abundant than CD8+ Tc cells in both GCNIS and GCNIS+ly. CD68+ macrophages, representing the second most abundant immune cell type in both sample categories, were markedly increased in GCNIS+ly compared to GCNIS. CD20cy+ B cells and CD11c+ DCs were mainly found in GCNIS+ly, but were rarely detected in GCNIS. CD25+ cells were observed in both GCNIS and GCNIS+ly, but FOXP3+ cells were mainly found in GCNIS+ly samples (Supplemental Fig. 1F). CXCR5+ cells were rarely observed in GCNIS and GCNIS+ly samples with no BCL6+ cells identified (Supplemental Fig. 1F).

The highest density of immune cell infiltration was found in seminoma and embryonic carcinoma samples, with CD3+ T cells, CD4+ Th and CD8+ Tc cells being more abundant in seminoma compared to embryonal carcinoma. The same applied for CD68+ macrophages, CD11c+ DCs, and CD20cy+ B cells. Importantly, a high density of cells expressing Treg-specific markers CD25 and FOXP3, or Tfh-specific markers CXCR5 and BCL6, were found mainly in seminoma (Supplemental Fig. 1F). BCL6+ cells were only found in seminomas containing follicular-like structures (FLS).

### Flow cytometry analysis revealed the highest number of immune cells in tumor central areas

Analysis of immune cells in tissue samples from different locations in both diseased and contralateral testes was employed to delineate the testicular immune cell composition of individual patients (Fig. 2A). Comprehensive analysis of samples obtained from 24 patients showed a marked variation in the frequency of immune cells at the different sampling sites of TGCT; the results illustrate the histopathological variation that exists within individual tumors. Despite the extensive phenotypic heterogeneity of the tissue samples based on histological assessment, flow cytometric analysis of immune cells unequivocally showed that CD3+ T cells were present at the significantly highest frequency in the tumor-central areas of seminoma (median 71.49%, range 19-89% of all cells) compared to all other locations in this and other TGCT categories (median 40.69%, range 6.36-77.32% of all cells) (Fig. 2B, Supplemental Table 4).

**Fig. 2:**
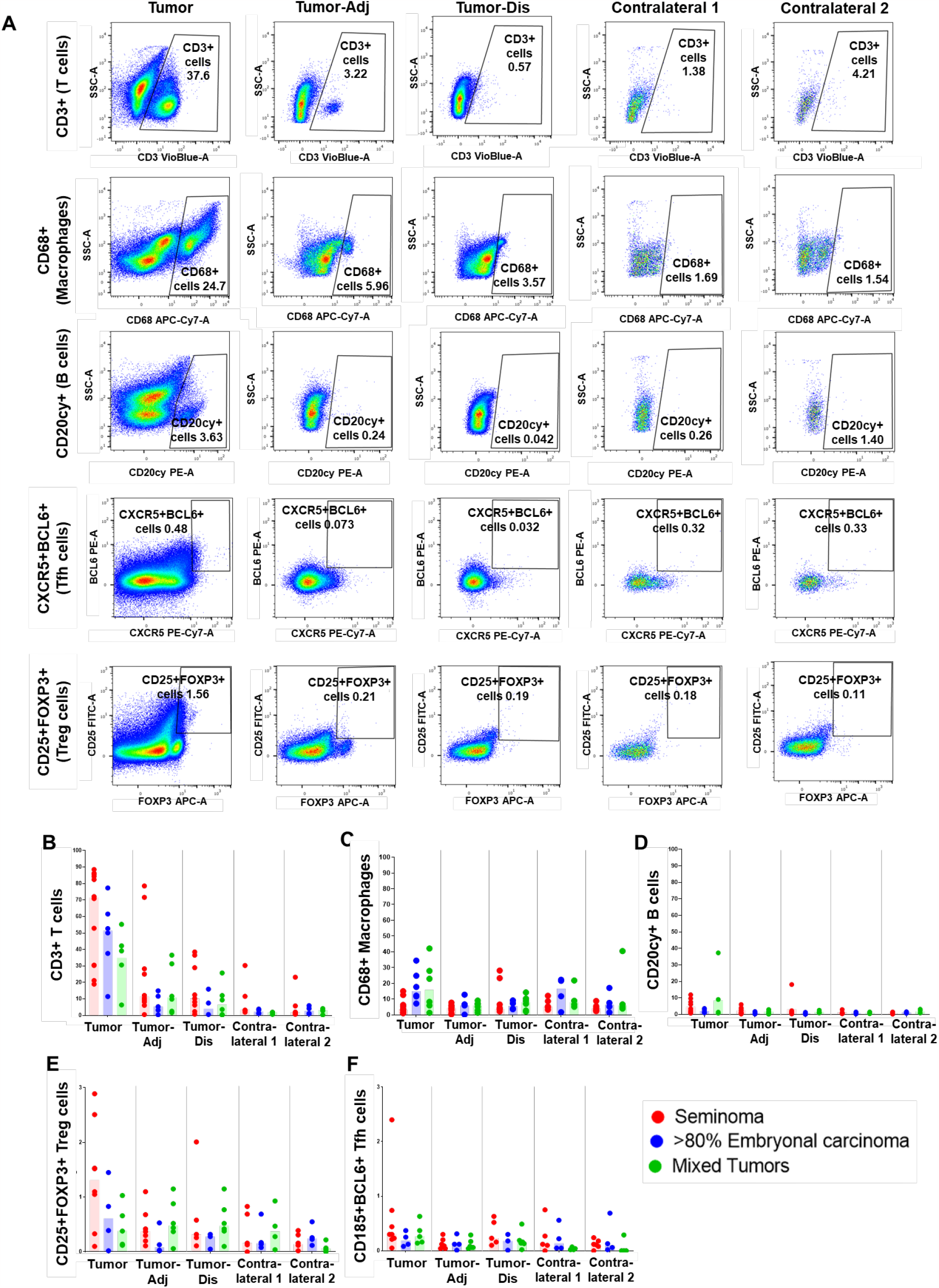
Flow cytometric analysis of human testicular immune cells. **A:** Representative analysis of immune cells in tissue samples from an individual patient with embryonal carcinoma, obtained from different localisations of both, diseased testis (Tumor; Tumor-adjacent [Tumor-Adj]; Tumor-distant [Tumor-Dis]) and contralateral testis ([Contralateral 1; Contralateral 2]). Percentage of positive cells with individual immune cell markers in total live cells is indicated. **B-D:** Overview of individual values measured for different immune cell markers in tumour-bearing and contralateral testes from n=24 patients [seminoma: Tumor n=12; Tumor-Adj n=10; Tumor-Dis n=9; Contralateral 1 n= 5; Contralateral 2 n= 6; embryonal carcinoma: Tumor n=6; Tumor-Adj n=5; Tumor-Dis n=4; Contralateral 1 n=4; Contralateral 2 n=4; mixed tumors: Tumor n=6; Tumor-Adj n=6; Tumor-Dis n= 6; Contralateral 1 n=4; Contralateral 2 n= 4]. **E-F**: Respective percentages of CD25+FOXP3+Treg and CXCR5+BCL6+ Tfh cells in total live cells [seminoma: Tumor n=8; Tumor-Adj n=7; Tumor-Dis n=5; Contralateral 1 n= 5; Contralateral 2 n= 6; embryonal carcinoma: Tumor n=4; Tumor-Adj n=4; Tumor-Dis n=3; Contralateral 1 n=4; Contralateral 2 n=4; mixed tumors: Tumor n=5; Tumor-Adj n=6; Tumor-Dis n= 6; Contralateral 1 n=4; Contralateral 2 n= 4].

Though numerically lower compared to in “Tumor” regions, significantly higher numbers of CD3+T cells were documented at “Tumor-Adj” and “Tumor-Dis” locations compared to the corresponding contralateral testes (Fig. 2B, Supplemental Table 4). Small proportions of CD3+T cells were also found in two of the four contralateral testis samples that exhibited normal spermatogenesis. Concomitant with the abundance of CD3+ T cells, the highest percentages of CD4+ Th cells and CD8+ Tc cells (17.03-60.69% and 0.56-43.98% of total live cells, with a median of 42.61 and 25.43%, respectively) were also detected in “Tumor” regions of seminoma (Supplemental Fig. 1C-D). CD68+ macrophages represented the second most frequent infiltrating immune cell in different sites of TGCT; they were the type most frequent in “Tumor” areas of embryonal carcinoma (Fig. 2C) and in contralateral testes, but were especially prominent in patients with embryonal carcinoma. CD20cy+ B cells were detectable at all sites in TGCT testes, but they were present in lower numbers compared to T cells and macrophages (Fig. 2D). The highest density of B cells was found within seminoma “Tumor” areas, followed by mixed tumors and embryonal carcinoma, but were rarely found in contralateral testes.

Flow cytometry showed a considerable abundance of CD25+FOXP3+ Treg cells in TGCT samples (Fig. 2E), with the highest percentage (0.10-2.89%, median of 1.31%) in seminoma “Tumor” areas, followed by embryonal carcinoma and mixed tumors (Fig. 2E). A small number of CD25+FOXP3+ Treg cells was also detected in the contralateral testes (Fig. 2E). CXCR5+BCL6+ Tfh cells were identified in all TGCT locations with the highest percentage (0.05-2.40%, median of 0.30%) in the seminoma “Tumor” sites (Fig. 2F). CXCR5+BCL6+ Tfh cells were rare in the “Tumor-Adj” and “Tumor-Dis” sites of the tumor-bearing testis (Fig. 2F) and in the contralateral testis (Fig. 2F).

### Single cell analysis of immune cells in normal human and TGCT reinforce abundance of T cells

Initial clustering analysis identified 46 individual clusters containing T cells, B cells, macrophages, plasma cells, endothelial cells, fibroblasts, germ cells, tumor cells, and undefined cells (Fig. 3A and Supplemental Fig. 2A) using both cell-specific signature markers (Fig. 3B and Supplemental Fig. 2B) and enrichment of differentially expressed genes (DEGs) in each cluster (Supplemental Fig. 2C). Results regarding cell numbers are presented as % of total analyzed cells after quality control. The sub-analysis of the datasets of normal testes (“donors”) included in the study revealed mostly germ cells (77.56%), accompanied by fibroblasts (2.90%) and endothelial cells (2.56%) and small proportions of immune cells, predominantly macrophages (1.82%) and few T cells (0.08%). No B cells were detected in human normal testis (Fig. 3A, C).

**Fig. 3:**
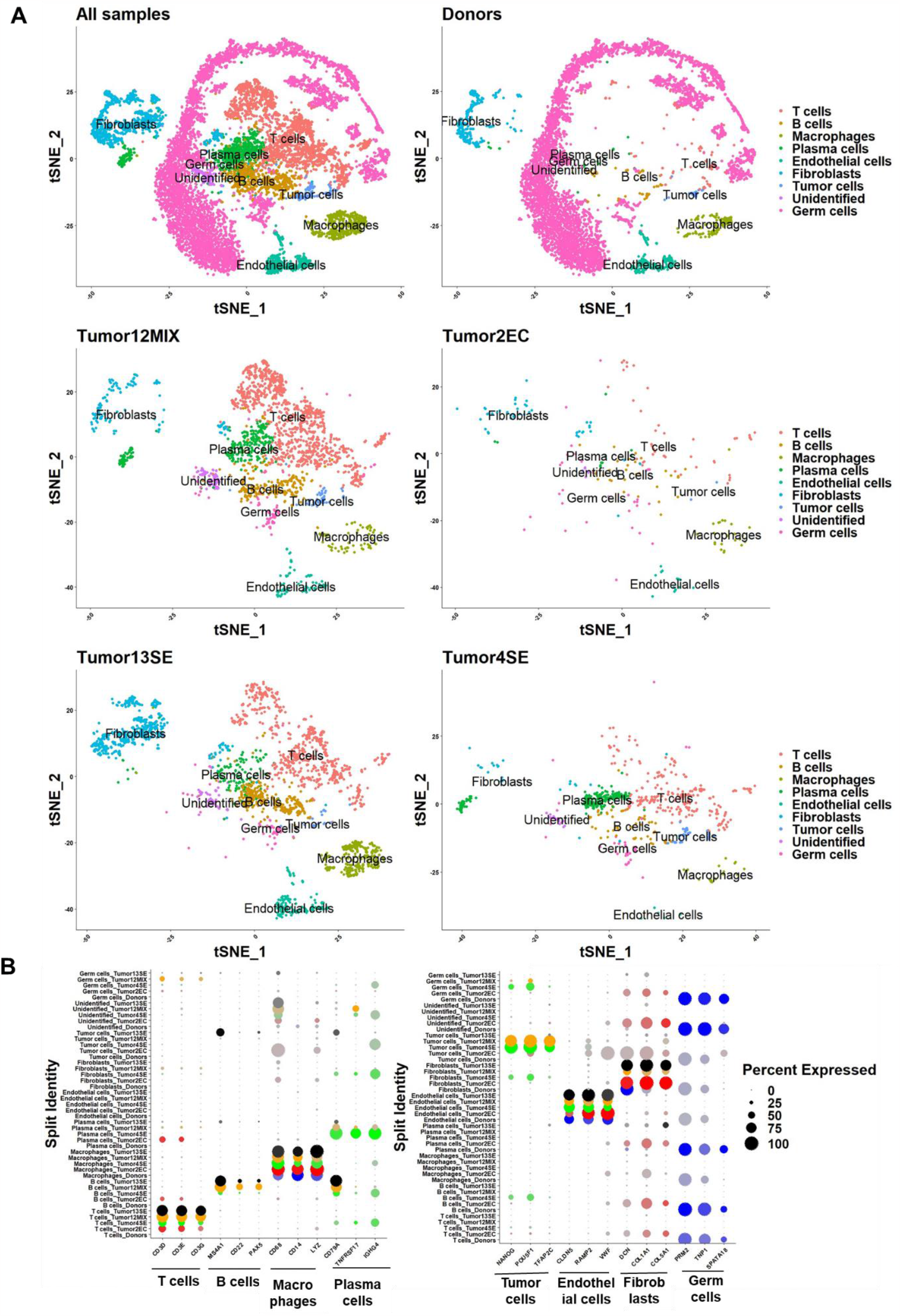
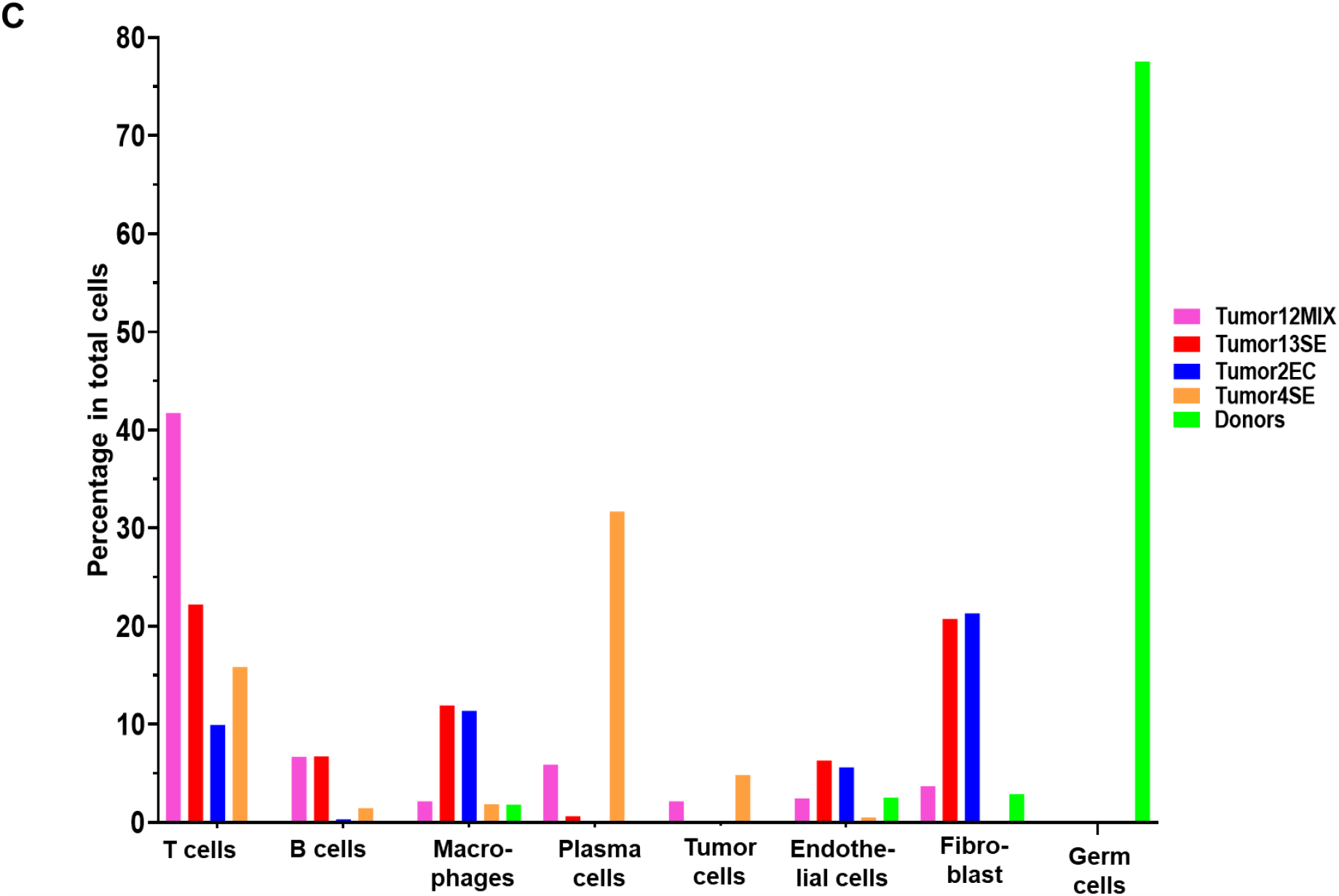
scRNA-seq analysis of normal human testis and TGCT identifies distinct cell populations, including immune cell subtypes. **A:** tSNE presentation of major cell types and associated clusters in healthy donor (n=3; pooled data) and TGCT samples (“Tumor12MIX” mixed TGCT, “Tumor13SE”: seminoma, “Tumor4SE”: seminoma, “Tumor2EC”: embryonal carcinoma). **B:** Dot Plot shows putative marker genes expressed by different cells in individual samples. Dot size encodes the percentage of cells within a cluster expressing the transcript, whereas the color intensity represents the average signal measured in expressing cells. **C**. The proportion of different cells among pooled healthy donor and individual tumor samples.

In agreement with the IHC and flow cytometry data, TGCT samples contained the highest numbers of T cells, B cells, and macrophages Fig. 3 A, C). T cells represented the major component of infiltrating immune cells (9.94-41.73%) (Fig. 3A, C), followed by macrophages (1.85-11.93%) and B cells (0.32-6.75%), respectively. TGCT samples also contained plasma cells (0.16-31.60%). However, the proportion of immune cell types significantly varied between tumor types: macrophages were the predominant immune cell type (11.37%) in embryonal carcinoma samples, whereas T cells accounted for 15.85-22.21% of the cells in seminoma and 41.73% in mixed tumors (Fig. 3A, C). In addition to immune cells, a considerable number of fibroblasts (0.0-20.72%), endothelial cells (0.5-6.33%), and tumor cells (0.08-4.83%) were found in TGCT (Fig. 3A, C).

Secondary clustering of T cells generated 17 subclusters, later identified by the canonical T cell markers (40) and enrichment of DEGs in each cluster (Fig. 4A, B and Supplemental Fig. 3A-C). In addition to cytotoxic T cells (6.00-33.67%), proliferating (activated) T cells (6.17-20%), CCR7+ (newly recruited) T cells (0.00-17.49), other CD4+ T cells (0.00-7.44%), and mixed T cells (9.98-28.89%), respectively, were present (Fig. 4C). Individual clusters representing Treg were detected in the secondary clustering analysis (0.00-16.80% of CD3+ T cells) (Fig. 4C), but no individual cluster for Tfh cells was identified. A newly developed analytical option in Seurat was employed to visualize and quantify the co-expression of CD4 and BCL6 to confirm the presence of Tfh in the available human testis samples (Fig. 4D). This advanced analysis identified Tfh cells (0.00-2.95% of CD3+ T cells) in the TGCT samples (Fig. 4E).

**Fig. 4:**
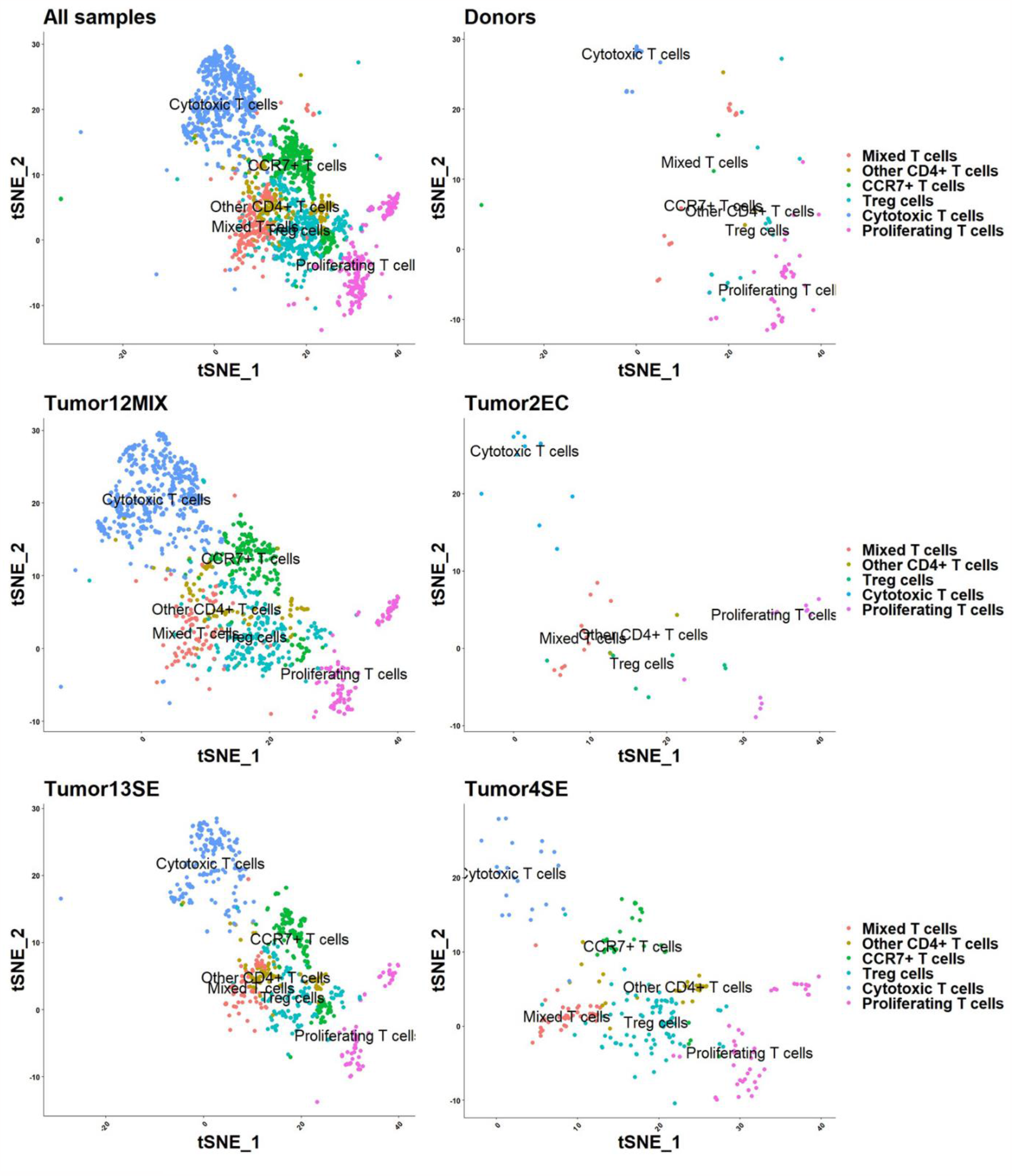
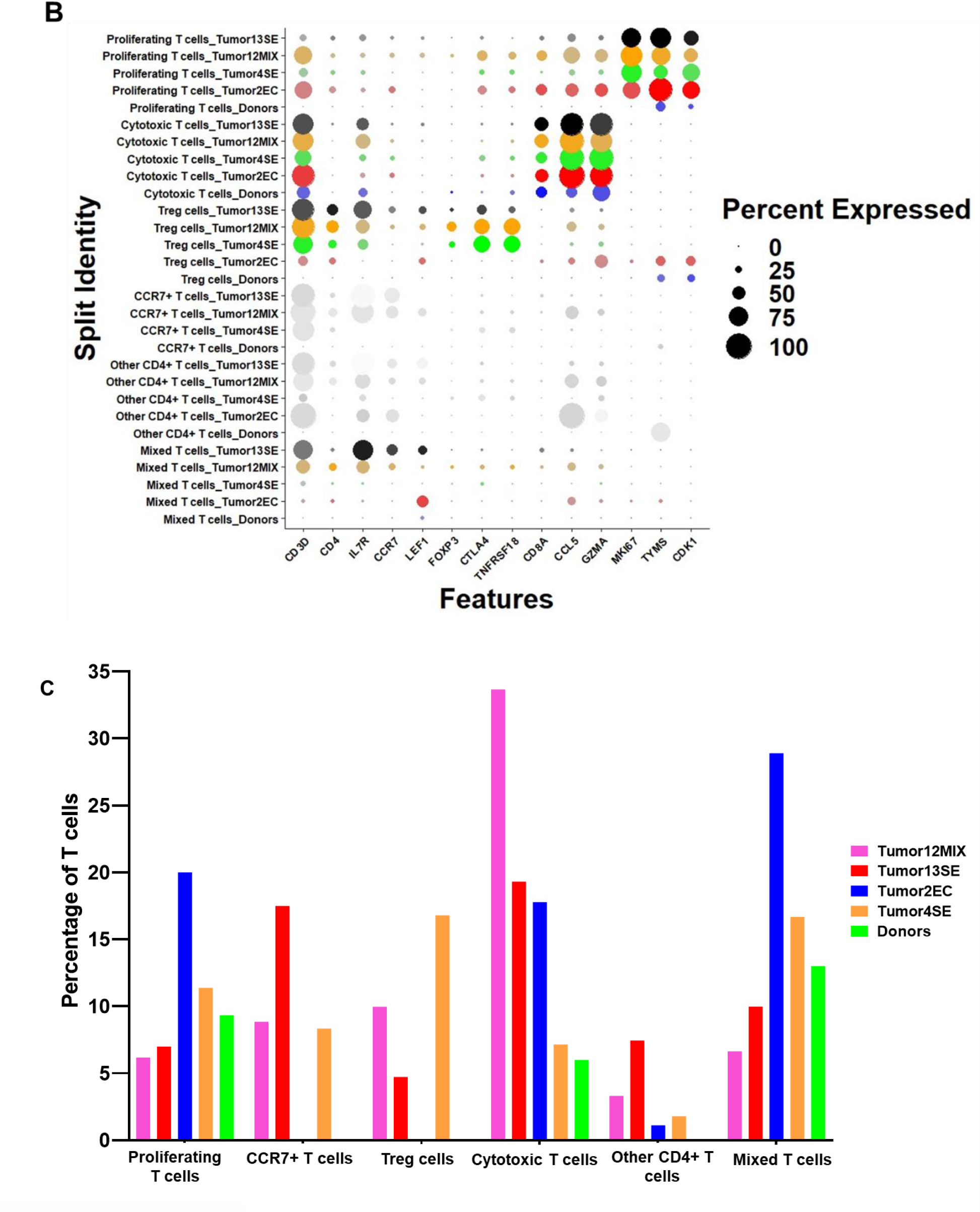
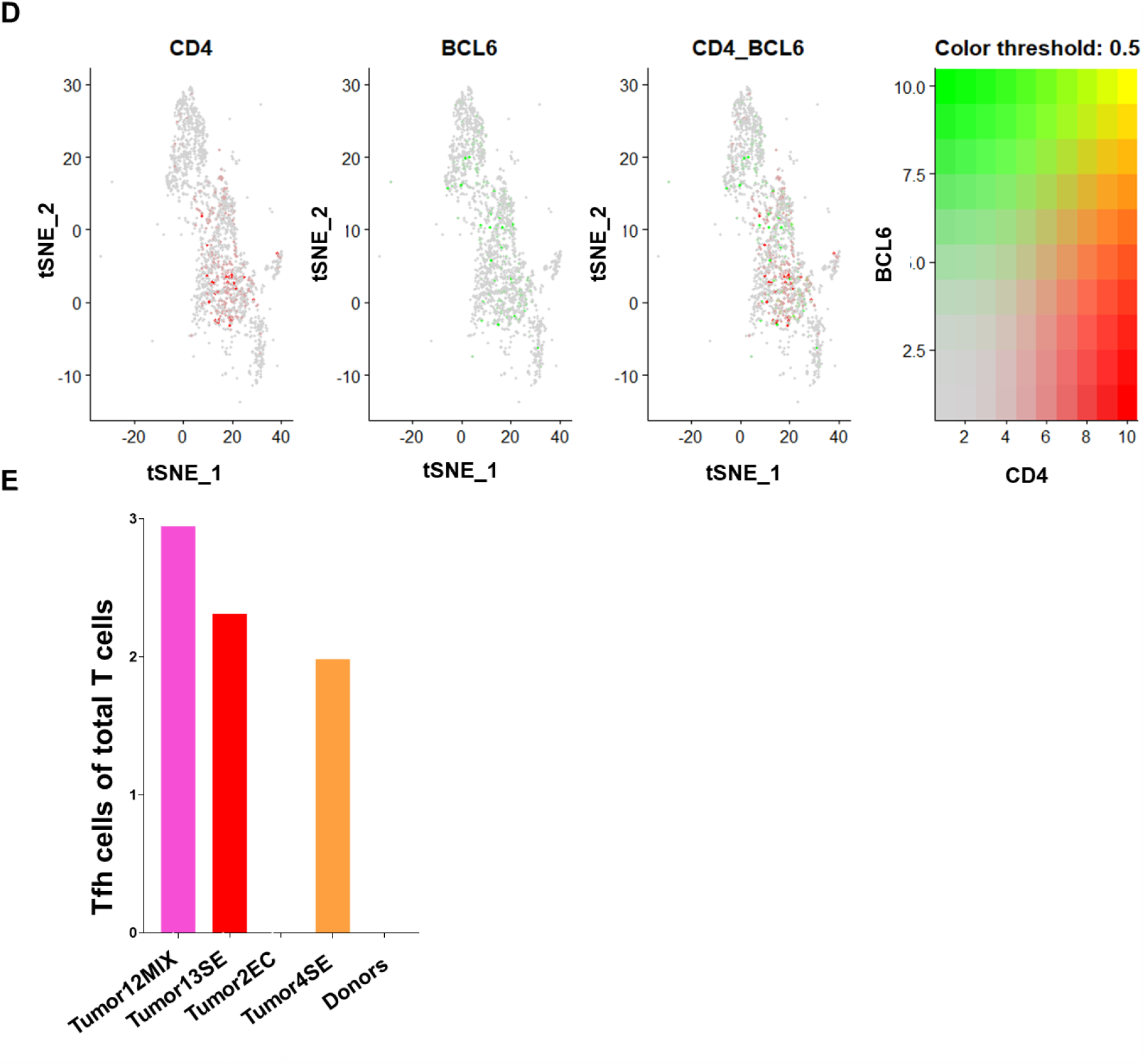
Analysis of T cells in normal human testis and TGCT. **A:** tSNE plot shows the clustering of T cells in donor (n=3; pooled data) and TGCT samples (Tumor12MIX: mixed TGCT, Tumor13SE: seminoma, Tumor4SE: seminoma, Tumor2EC: embryonal carcinoma). **B:** Dot plot shows expression levels of immune typing markers in different T cells clusters in a pooled analysis of all studied samples. The size of the dot encodes the percentage of cells within a cluster expressing the gene, whereas the color intensity encodes the average expression level of expressing cells. **C:** Bar plot shows the proportion of different T cell subtypes among donors and TGCT. **D:** Co-expression of Tfh cell markers (CD4 and BCL6) in total T cell population. **E:** Proportion of Tfh cells in individual samples.

## DISCUSSION

Applying flow cytometry in tissue specimens from TGCT-bearing and unaffected contralateral testes, we were able to reinforce the results obtained from corresponding immunohistochemical evaluation and previous results from our group (8). In 96% of TGCT patients, the proportion of CD4+ cells exceeded that of CD8+ TIL, with their numbers gradually decreasing from Tumor to peripheral areas, and significantly fewer T cells in tumor-free, contralateral testes. In our TGCT cohort, CD4+ cells were more abundant in seminoma compared to embryonal carcinoma and other TGCT entities, while Siska et al (9) found no difference in CD4+ and CD8+ T-cell infiltration in seminomas vs. non-seminomas. This could be due to the expression of CD4 by macrophages (46), so CD4+ cells detected by IHC in our study could also include interstitial macrophages. However, the IHC results in our study were supported by flow cytometry, because CD4+ and CD8+ cells were measured as proportion of CD3+ cells, and hence only CD4+ T cells would be counted in this latter analysis (data not shown).

Both IHC and flow cytometry results were confirmed in independent samples using scRNA-seq, which also revealed a marked heterogeneity among TGCT patients and proved T cells to be the most abundant immune cell in TGCT, while macrophages were predominant in normal testes. Studies addressing the T cell compartment in the normal and diseased human testis including TGCT in detail are rare. In other cancers, Treg heavily infiltrate the TME and are critically associated with poor prognosis in lung, gastric, ovarian cancer, melanoma etc. (29-31). Despite sample heterogeneity, flow cytometric analysis revealed a significant increase of CD25+FOXP3+ Treg cells in seminoma, compared to non-neoplastic testicular tissue, suggesting an important role in TGCT development. While corresponding IHC confirmed the results of flow cytometry, other IHC-based studies revealed decreased numbers of FOXP3+ cells in testes with focal inflammatory lesions as well as GCNIS (7, 47), which is most likely due to the different panel of markers and samples used.

Treg exhibit their immunosuppressive activities by interacting with different immune cells including CD4+ Th, CD8+ Tc, B cells, DC, and macrophages (29-31). The co-localization of Tregs with these other immune cells suggests they are interacting in testicular neoplasia. The profoundly different immune landscape in neoplastic testes was validated by our scRNAseq data. In addition to CD8+ Tc cells, proliferating (activated) T cells, CCR7+ (newly recruited) T cells, and Treg cells could be identified among CD4+ T cells at the transcriptome level in TGCT, but not in normal testes. In line with the high inter-patient variation observed in our study, differential expression of *CD3D, CD3E*, and genes associated with TIL senescence were reported in two seminoma subtypes (48, 49).

Our previous work showed that, along with pro-inflammatory (IL-1β, IL-6, TNF-α) and Th1-related cytokines (IL-2, IFN-γ), transcripts encoding Treg related cytokines TGF-β and IL-10 were significantly higher in TGCT compared to NSP or non-neoplastic inflammation (HYP+ly) (8). Despite the high consumption of IL-2 by Treg and their ability to down-regulate or directly destroy effector T cells (29-31, 50), our results clearly showed other CD4+ and CD8+ T cell subsets were not depleted from the TME. In contrast, Siska et al. reported decreased pan-T cell signatures associated with advanced TGCT stage, while Treg signatures were increased (9).

To the best of our knowledge, this is the first report demonstrating the presence of Tfh cells in the human testis. Tfh cells can be positively (e.g., breast cancer, colon cancer, pancreatic ductal adenocarcinoma) or negatively (e.g., hepatocellular carcinoma) associated with the long-term survival of patients through releasing CXCL-13, PD-1, CXCR-5, ICOS and BCL6 as well as interacting with B cells and CD8+ Tc cells (32-35, 51). IHC revealed that Tfh cells are most abundant in seminoma and localized within FLS, which also contain B cells, CD4+ Th cells, and follicular DC. Consistent with previous reports (8, 19), tumor-infiltrating B cells were exclusively found in TGCT but were virtually absent in normal testis tissue in our analysis. The presence of Tfh and clustering of B cells in FLS are both in line with increased transcript levels of supporting chemokines in testicular germ cell neoplasia, i.e. CXCL-13 and CCL-5 (8). In the TME of different cancers, Tfh cells have been identified as the main source of IL-21, another B cell-supporting cytokine (34). Additionally, TGCT are associated with high IL-6 expression, with implications for differentiation, proliferation and activation of immune cells including B cells, DC, and T cell subtypes (8, 22, 52). IL-6 can promote malignant cell proliferation, angiogenesis, and metastasis in other cancers (53). Accordingly, Nestler et al. reported that IL-6 signalling is the most significantly upregulated immune response pathway in metastatic versus non-metastatatic seminoma, thus impacting patients’ prognosis (22).

Our results provide evidence that recruitment of non-resident immune cells contributes to the unique TME in TGCT. Although CD8+ Tc could be identified in all cohorts and experimental approaches, it remains to be elucidated whether these cells can exert TGCT-specific cytotoxicity, or rather have regulatory or ‘bystander’ functions, comparable to CD4+ subsets (7). Similarly, the specificity of clonally-expanding B cells in TGCT remains to be unravelled (19). It is a matter of debate whether anti-inflammatory priming and/or dysfunction of TIL might enable neoplastic germ cells to escape from immune surveillance, and thereby support tumor development and progression. Clinical trials targeting immune checkpoint molecules such as PD-1 for the treatment of TGCT mainly failed due, probably to our incomplete understanding of the TGCT immune microenvironment (36, 49, 54), but IL-6 has been considered as a potential immunotherapeutic target in seminoma (8, 22). In other cancers, IL-21 blockade was able to drastically reduce B cell activation induced by co-administration of anti-PD-1 and anti-CTLA-4 therapy, highlighting the importance of Tfh cell-secreted effector molecules in cancer immunotherapy (55).

The role of pro-inflammatory cytokines in the TME is supported by an in vitro co-culture model using the human seminoma cell line TCam-2 and immune cell fractions, where TCam-2 cells induce immune cells activation and generate a strong pro-inflammatory milieu through producing IL-2, IL-6 and TNFα (52, 56, 57). In addition, TCam-2 showed an immediate increase in pro-inflammatory cytokine mRNA levels (IL-1β, IL-6, TNF-α, etc.) and is capable to produce IL-6 after direct contact with PBMC (52).

The development of personalized oncological medicine in TGCT is hampered by inter-individual variation and intra-tumoral heterogeneity which was also revealed in the present study. The marked regional immune-related gene expression differences, correspond to inter- and intra-tumoral heterogeneity and indicate the importance of the immune landscape in TGCT progression and metastatic spread (22). Similarly, our flow cytometric analysis showed a clear difference in the composition of immune cells at different locations of tumor-bearing and contralateral testis tissues with highest numbers in tumor-central sites of TGCT. Rare Treg and Tfh cells were also most prominent at Tumor sites compared to other locations, indicating their important involvement in TGCT.

Limitations of our study are related to restricted access to human testis tissue and the obvious inter-individual and intra-sample heterogeneity. Moreover, the small amount of each sample available for research risks the loss of rare immune cell subsets during preparation of single cells. The combination of experimental approaches applied here does not give access to spatial cell-cell interactions that might reveal functional interactions, and downstream analysis of transcriptome data from different immune cell populations in normal human testis and TGCT would require additional samples, and is thus beyond the scope of this study.

In conclusion, these findings data add depth to knowledge of the complex immune environment of TGCT, beyond conventional histopathology. Despite high inter-individual variation and sample heterogeneity, all experimental approaches showed a consistent immune cell pattern with a major increase of immune cells in central areas of TGCT compared to normal testis. The predominance of resident macrophages under physiological conditions is shifted to tumor-infiltrating T cells with an increased proportion of rare T cell subtypes, particularly Treg and Tfh, providing first evidence of Treg and Tfh involvement in TGCT biology. Further studies are needed to functionally characterize the testicular immune cell landscape, thus improve our understanding of immune surveillance in TCGT development and progression, and define potential targets for personalized immunotherapy.

## ACKNOWLEDGEMENTS

The authors greatly appreciate Ms. Alexandra Hax, Ms. Kerstin Wilhelm and Ms. Tania Bloch for their competent technical support. Many thanks to the Monash Bioinformatics Platform for their bioinformatics support.

## AUTHOR CONTRIBUTION

DF, HC-S, KL, MH, BL designed the study. JG, XW, XN and JHH provided scRNA-seq data. RI, JH and MF conceptually contributed to the work and performed pathological evaluation and IHC. RI performed flow cytometric analysis and scRNA-seq data analysis. BN helped with the bioinformatics. CP, MF contributed to the design and optimization of the flow cytometry panels. SI, KH were conceptually involved in the work. SK, FD, AP, FW, HC-S provided all specimens. RI, HC-S, and DF prepared the manuscript. RI, MH, BL, JG, HC-S, DF, and KL made the final critical revision of the manuscript. All authors read and approved the final version of the manuscript.

## ETHICS APPROVAL AND CONSENT TO PARTICIPATE

This study was approved by the ethics committee of the Medical Faculty of the Justus Liebig University Giessen; Ref. No. 26/11; 152/16 and the University of Utah Andrology laboratory consented for research (IRB approved protocol #00075836).

**Supplemental Fig. 1: Analysis of immune cells of human testis samples by flow cytometry and IHC. A:** Schematic illustration of sample collection sites during surgery of TGCT (Tumor: within tumor; Tumor-Adj: adjacent to tumor; Tumor-Dis: distant from tumor; Contralateral 1: upper pole of the contralateral testis, Contralateral 2: lower pole of the contralateral testis), and workflow of flowcytometry. **B:** Flowcytometry using antibody panel-1 (CD45, CD20cy, CD68, CD3, CD4, CD8) and respective gating strategy to analyse different testicular immune cells. **C-D:** presence of CD4+ and CD8+ T cells in total live cells, CD4+ T cells are more frequent in “Tu” sites of seminoma samples than in other localisations and compared to CD8+ T cells. **E:** Antibody panel-2 (CD45, CD3, CD4, CD25, FOXP3, CXCR5, BCL6) and respective gating strategy to analyse Treg and Tfh cells. **F:** Corresponding results of IHC analysis (seminoma, n=26 and embryonal carcinoma, n=10) compared to NSP (n=10), HYP+LY (n=12), GCNIS (n=14), GCNIS+LY (n=15) showing that Treg and Tfh cells are mostly found in TGCT samples, especially in SE. White bar in top left-hand panel indicates 100μm; all images at the same magnification.

**Supplemental fig. 2: Clustering information of different cell landscape in normal testis and TGCT. A:** tSNE presentation of major cell types and associated clusters in all samples. **B:** tSNE plots and Violin plots show the expression of selected markers for each cell type throughout 46 clusters. **C**. Heatmap of the top 5 significantly differentially expressed genes in each cell type analyzed.

**Supplemental fig. 3: Clustering information of different T cell landscape in normal testis and TGCT. A:** The tSNE plots align the T cell clusters. **B:** tSNE plots and Violin plots show the expression of selected marker for different T cell subtypes. **C:** Heatmap shows the top 5 differentially expressed genes by various T cell clusters in the studied samples.

## Supplemental Tables

**Supplemental Table 1:** List of antibodies used for IHC and their working conditions.

**Supplemental Table 2: List of the antibodies with their conjugated dye used for Flow cytometric analysis and their working conditions**. All antibodies were purchased from Miltenyi Biotec (except FOXP3, Biolegend). Intercellular targeted antibodies are marked with blue color.

**Supplemental Table 3: Number of cells in the scRNA-seq data sets analyzed in the current study**.

