## Supplemental Figures for "T cells in testicular germ cell tumors: new evidence of fundamental contributions by rare subsets"

### Supplemental Fig. 1

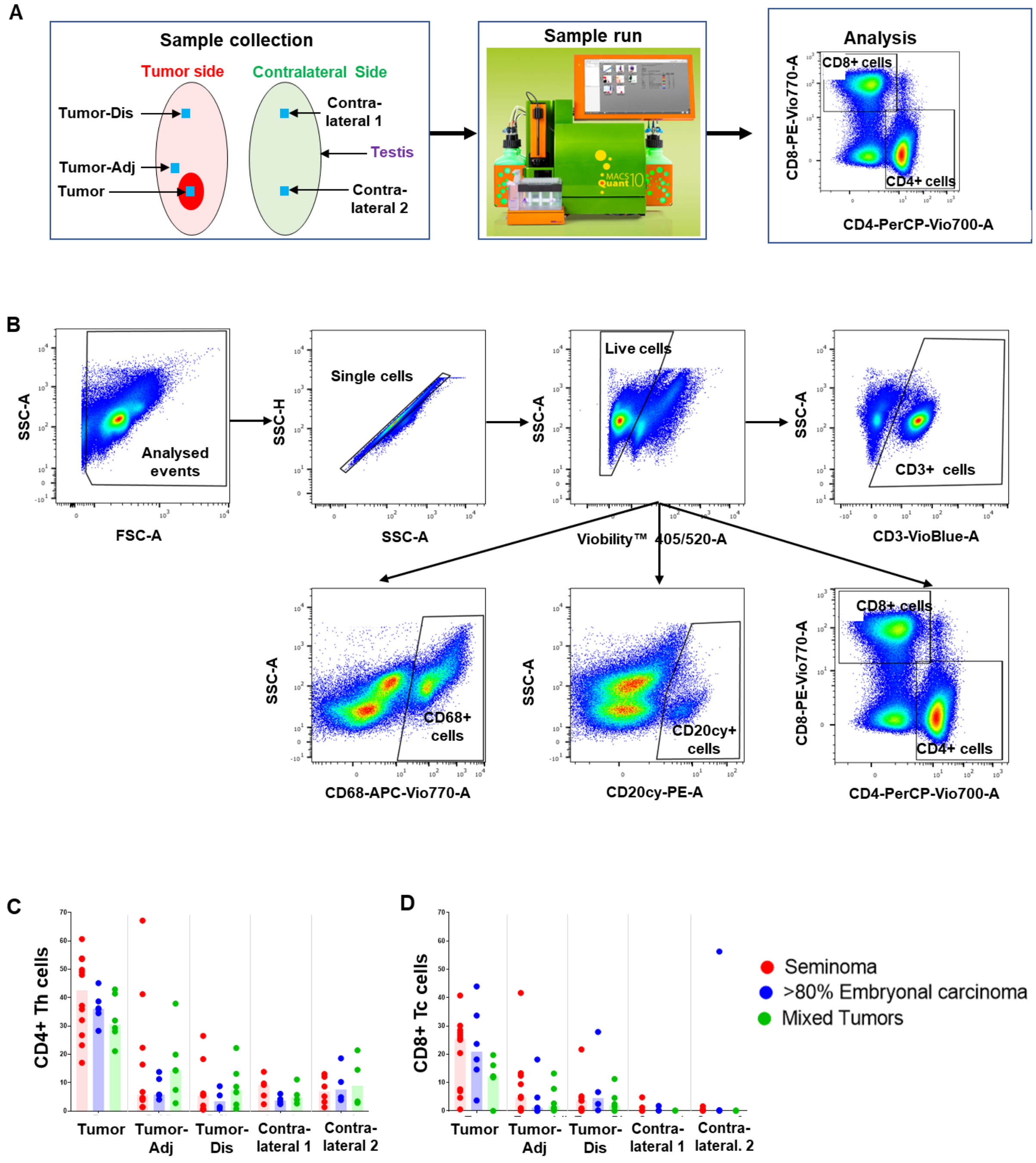


Supplemental Fig. 1 Cont.

**E**

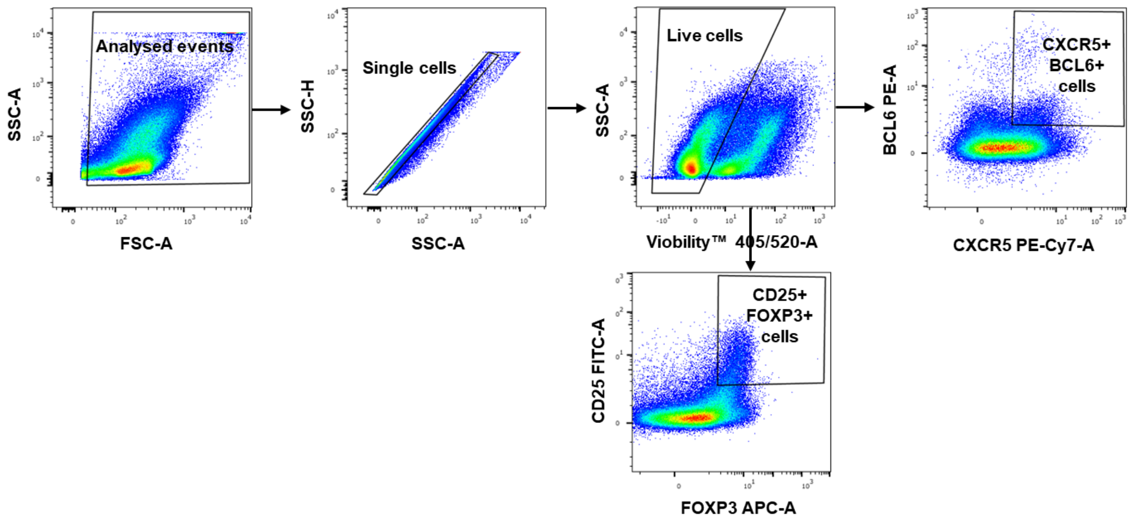
Analysed events
SSC-A
FSC-A
Single cells
SSC-H
SSC-A
Live cells
SSC-A
Viobility™ 405/520-A
CXCR5+
BCL6+
cells
BCL6 PE-A
CXCR5 PE-Cy7-A
CD25+
FOXP3+
cells
CD25 FITC-A
FOXP3 APC-A

**F**

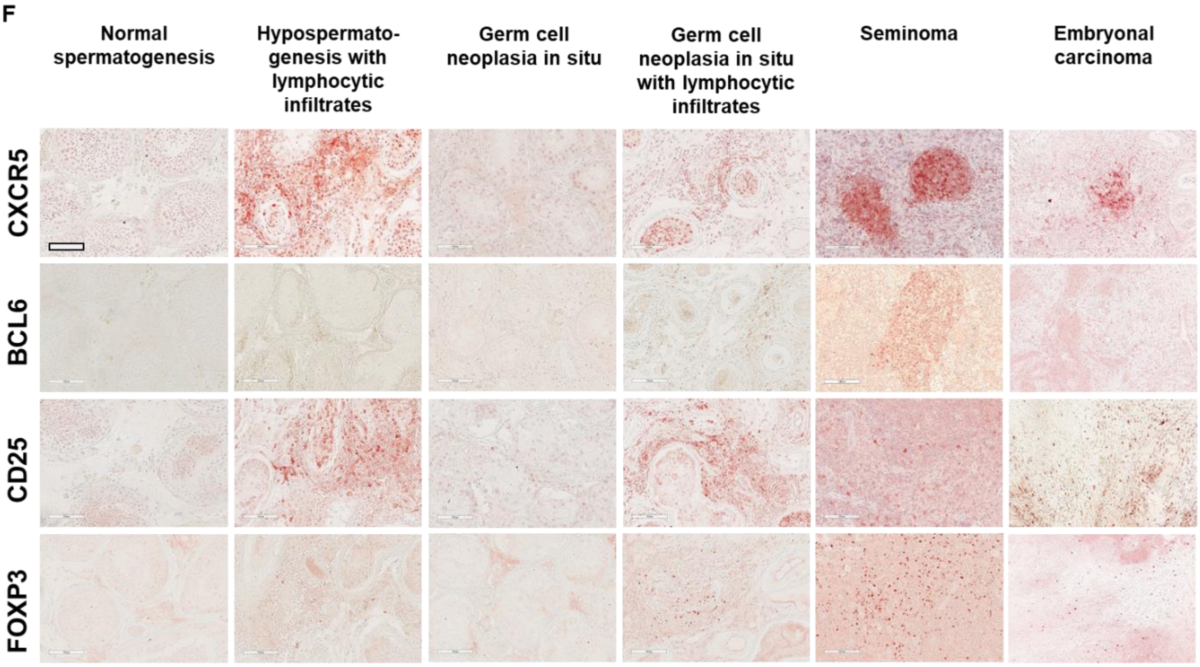
Normal spermatogenesis
Hypospermato-genesis with lymphocytic infiltrates
Germ cell neoplasia in situ
Germ cell neoplasia in situ with lymphocytic infiltrates
Seminoma
Embryonal carcinoma
CXCR5
BCL6
CD25
FOXP3

**Supplemental Fig. 2**

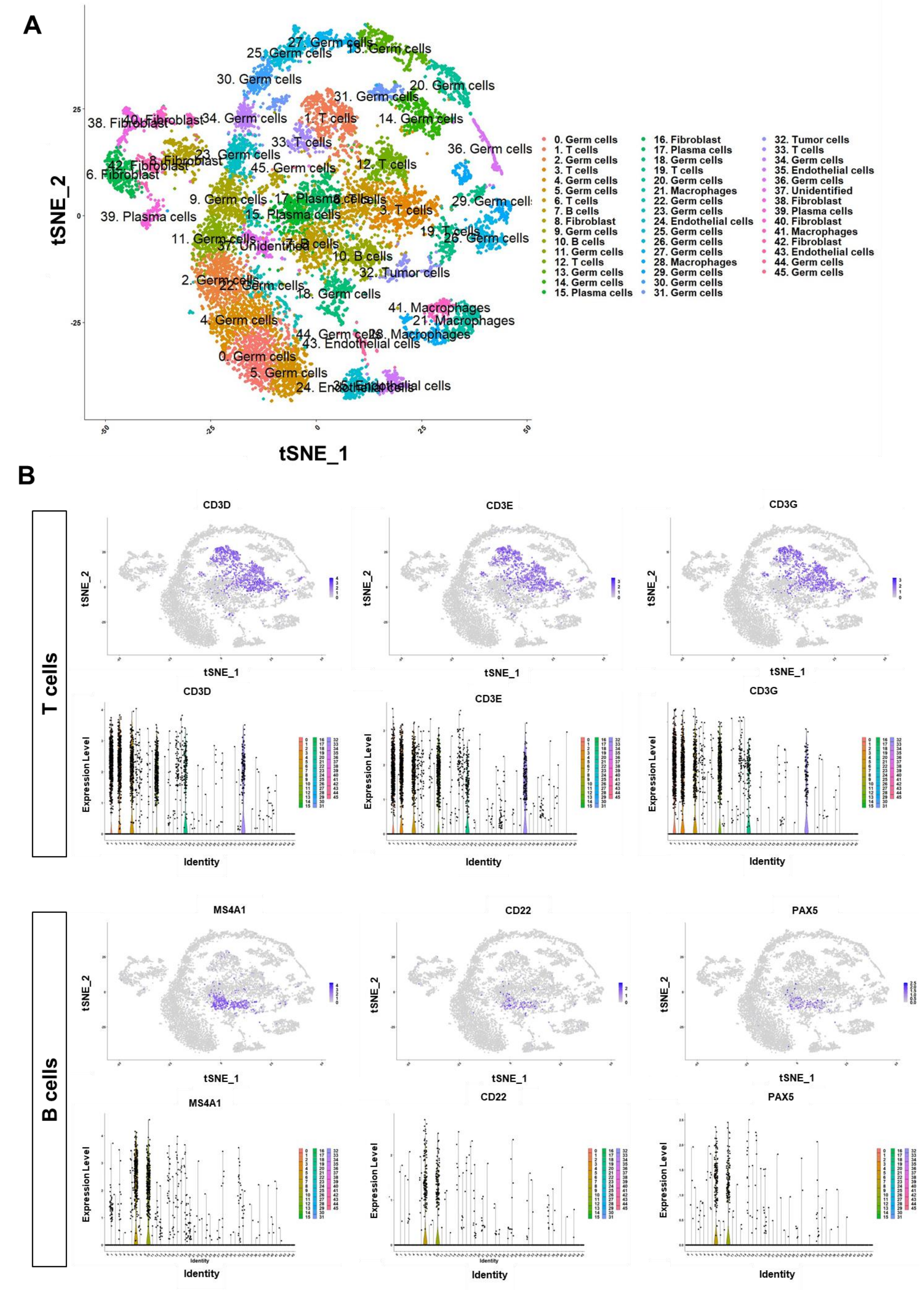

B. Cont.

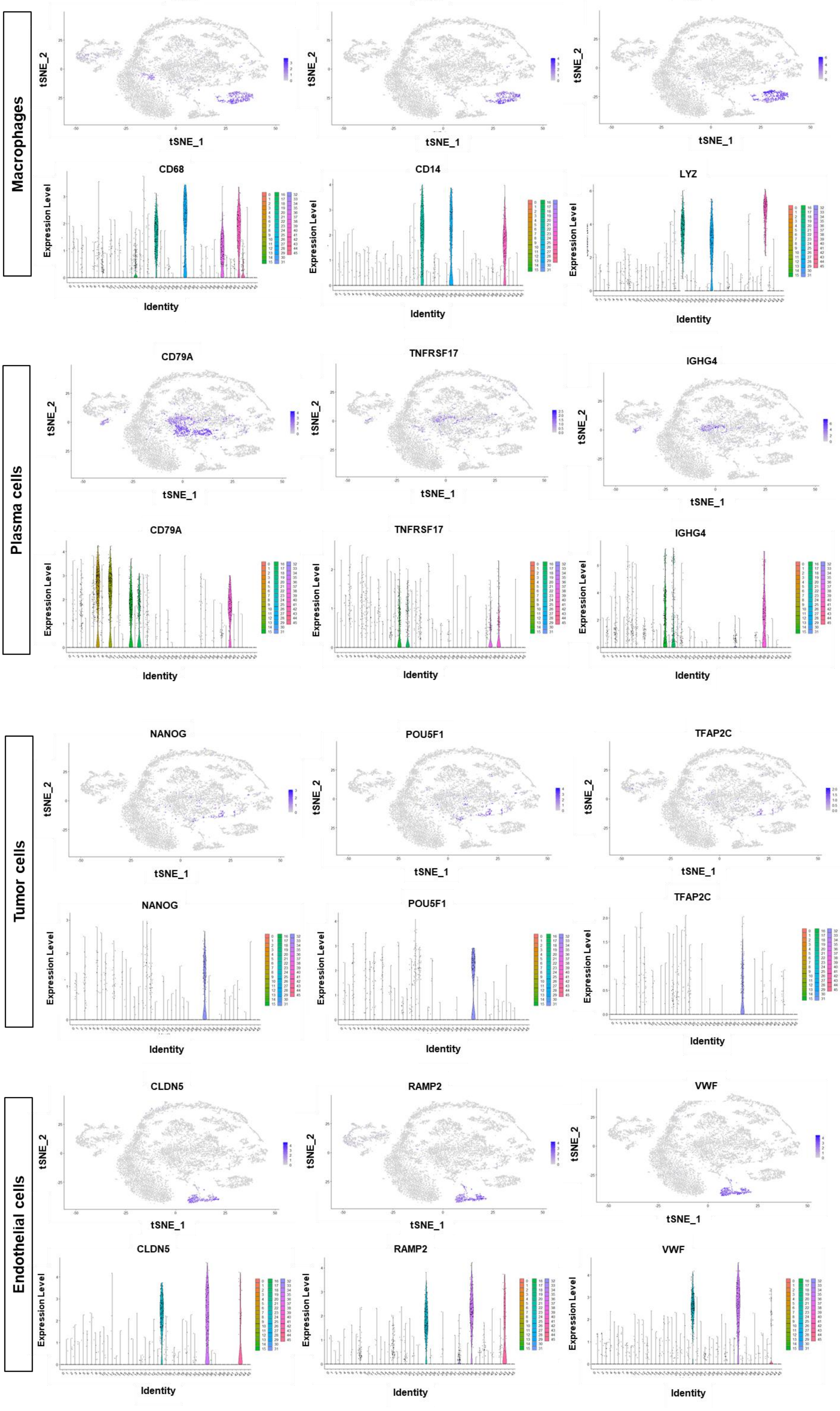
Macrophages
CD68
CD14
LYZ
tSNE_2
tSNE_1
Expression Level
Identity
Plasma cells
CD79A
TNFRSF17
IGHG4
Tumor cells
NANOG
POU5F1
TFAP2C
Endothelial cells
CLDN5
RAMP2
VWF

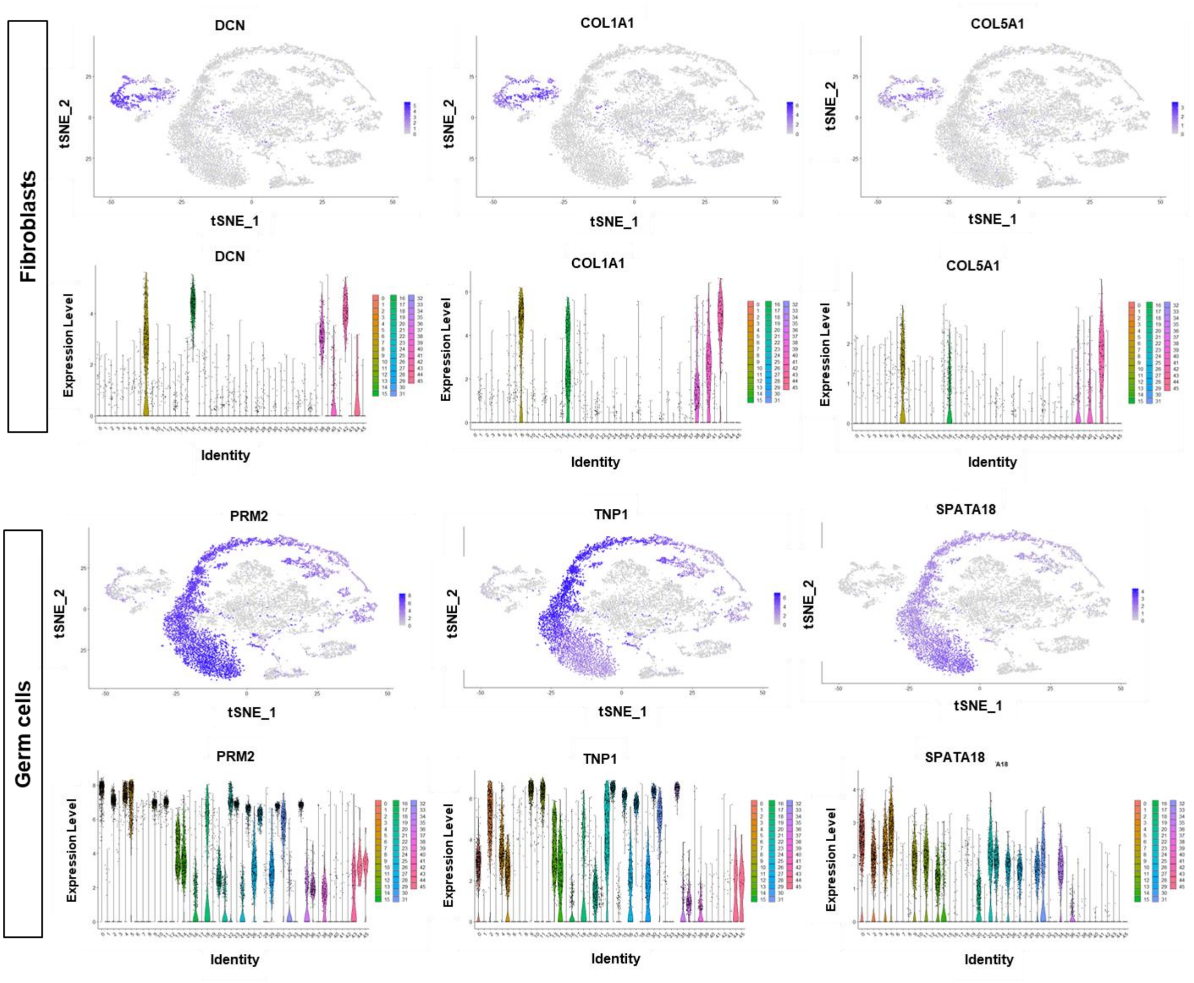
Fibroblasts
DCN
COL1A1
COL5A1
tSNE_2
tSNE_1
Expression Level
Identity
Germ cells
PRM2
TNP1
SPATA18

C

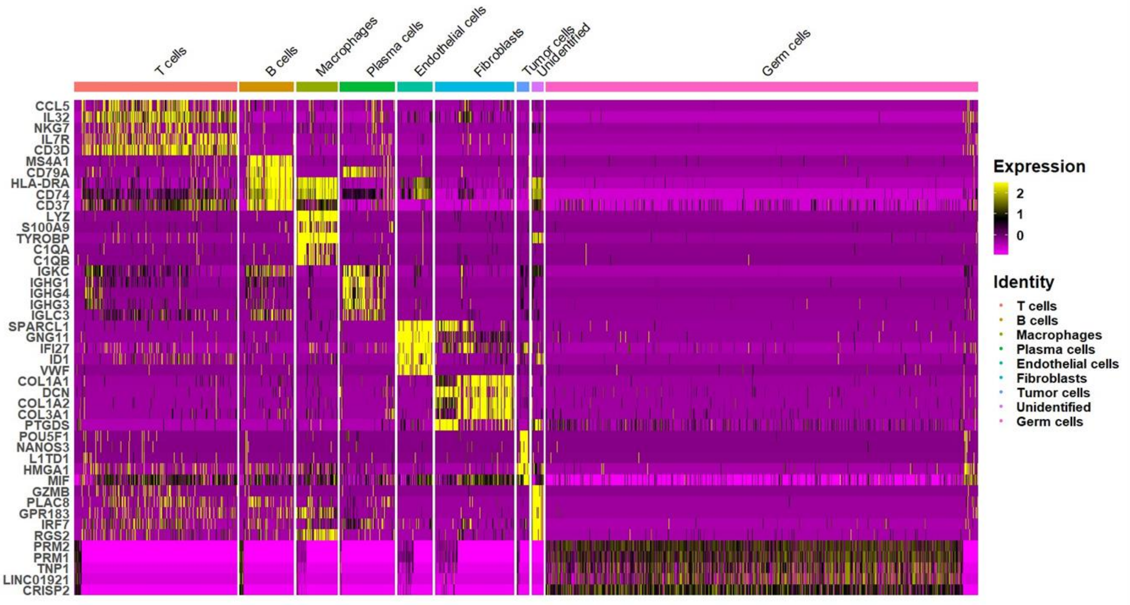
T cells
B cells
Macrophages
Plasma cells
Endothelial cells
Fibroblasts
Tumor cells
Unidentified
Germ cells
CCL5
IL32
NKG7
IL7R
CD3D
MS4A1
CD79A
HLA-DRA
CD74
CD37
LYZ
S100A9
TYROBP
C1QA
C1QB
IGKC
IGHG1
IGHG4
IGHG3
IGLC3
SPARCL1
GNG11
IFI27
ID1
VWF
COL1A1
DCN
COL1A2
COL3A1
PTGDS
POU5F1
NANOS3
L1TD1
HMGA1
MIF
GZMB
PLAC8
GPR183
IRF7
RGS2
PRM2
PRM1
TNP1
LINC01921
CRISP2
Expression
2
1
0
Identity
T cells
B cells
Macrophages
Plasma cells
Endothelial cells
Fibroblasts
Tumor cells
Unidentified
Germ cells

### Supplemental Fig. 3

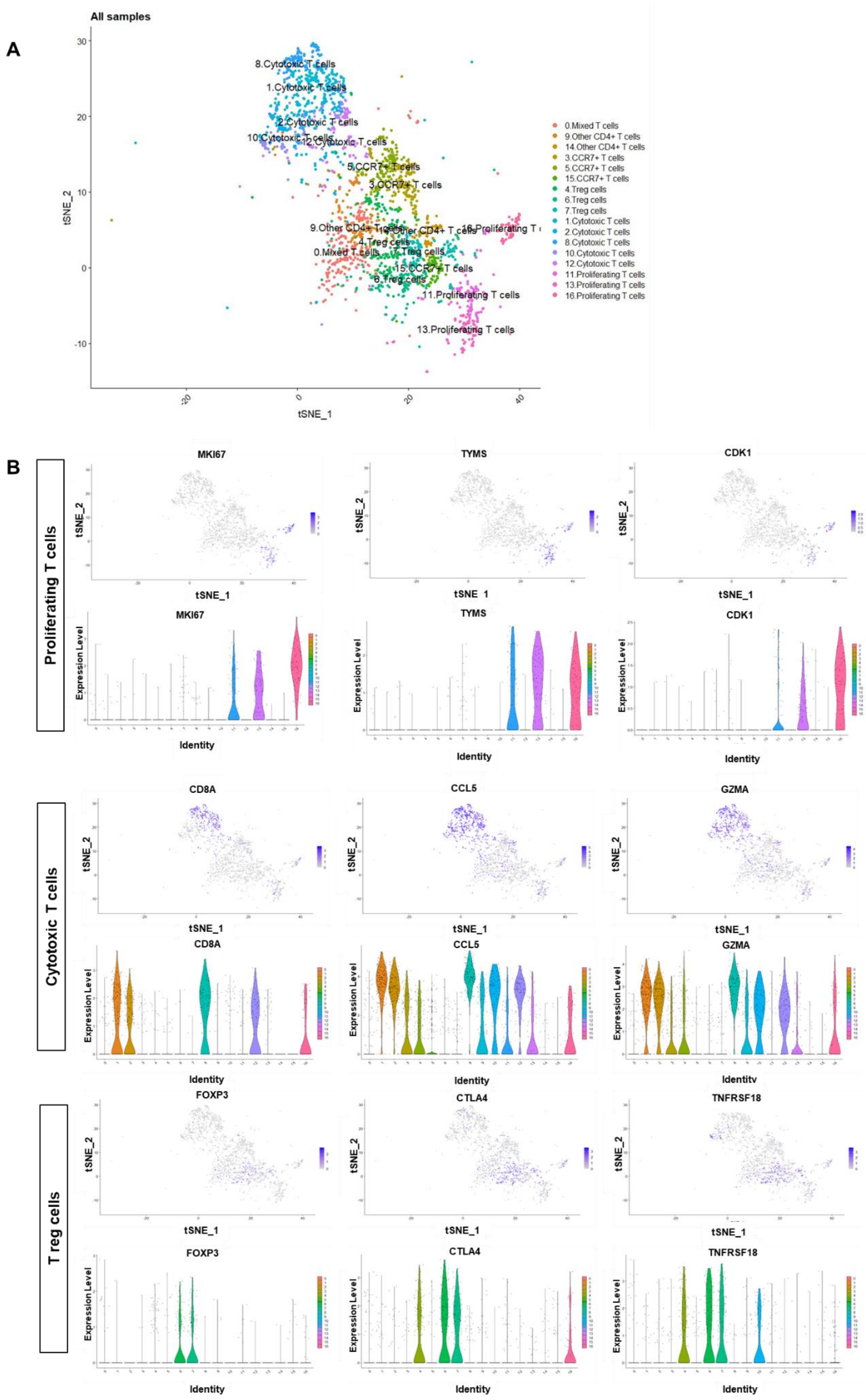

**B cont.**

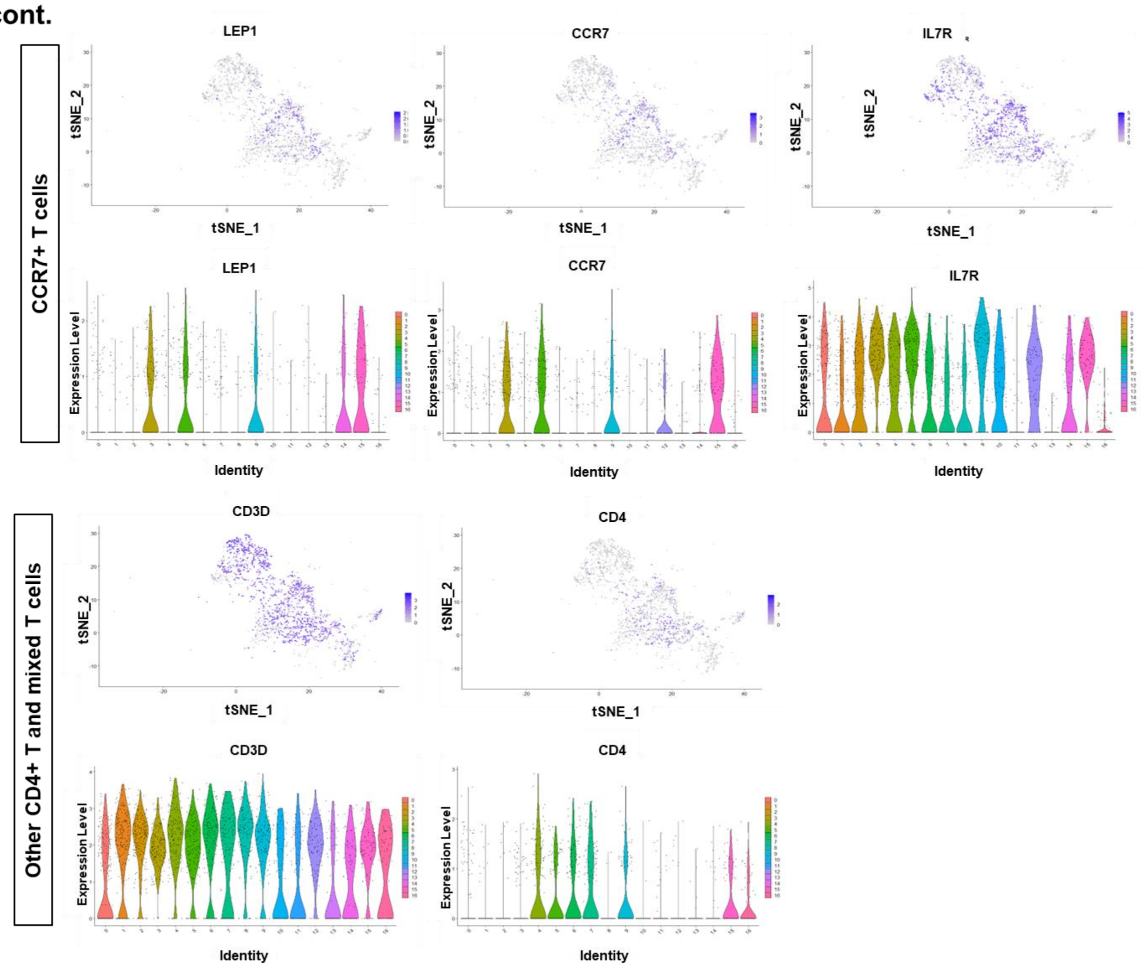
CCR7+ T cells
LEP1
CCR7
IL7R
tSNE_2
tSNE_1
Expression Level
Identity
Other CD4+ T and mixed T cells
CD3D
CD4

**C**

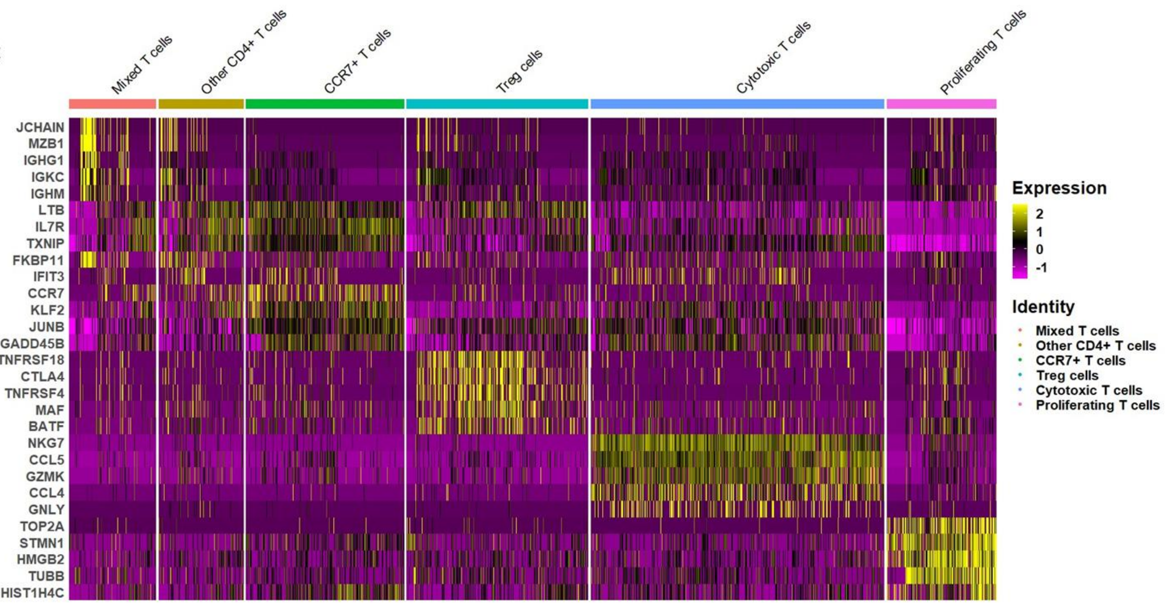
Mixed T cells
Other CD4+ T cells
CCR7+ T cells
Treg cells
Cytotoxic T cells
Proliferating T cells
JCHAIN
MZB1
IGHG1
IGKC
IGHM
LTB
IL7R
TXNIP
FKBP11
IFIT3
CCR7
KLF2
JUNB
GADD45B
TNFRSF18
CTLA4
TNFRSF4
MAF
BATF
NKG7
CCL5
GZMK
CCL4
GNLY
TOP2A
STMN1
HMGB2
TUBB
HIST1H4C
Expression
2
1
0
-1
Identity
Mixed T cells
Other CD4+ T cells
CCR7+ T cells
Treg cells
Cytotoxic T cells
Proliferating T cells
