## Supplemental Tables for "T cells in testicular germ cell tumors: new evidence of fundamental contributions by rare subsets"

**Supplemental Table 1:** List of antibodies used for IHC and their working conditions.

| Primary antibody | Specificty | Supplier and Catalogue no | Dilution | Staining |
|---|---|---|---|---|
| Polyclonal rabbit anti-human CD3 | Pan T cells | DAKO, A0452 | 1:100 | AEC |
| Monoclonal mouse anti-human CD20cy | B cells | DAKO, M0755 | 1:100 | AEC |
| Monoclonal mouse anti-human CD68 | Macrophages | DAKO, M0876 | 1:100 | NovaRED |
| Monoclonal mouse anti-human CD11c | Dendritic cells | Novocastra, NCL-L-CD11c-563 | 1:100 | NovaRED |
| Monoclonal rabbit anti-human CD4 | T helper (Th) cells | Abcam, ab133616 | 1:100 | AEC |
| Monoclonal mouse anti-human CD8a | Cytotoxic T (Tc) cells | eBioscience, Product: 14-0008-82 | 1:250 | AEC |
| Polyclonal rabbit anti-human IL2RA (CD25) | Treg | Sigma, HPA054622 | 1:500 | AEC |
| Monoclonal mouse anti-human FOXP3 | Treg | eBioscience, Product: 14-4777-80 | 1:100 | AEC |
| Polyclonal rabbit anti-human CXCR5 | Tfh | Sigma, HPA042432 | 1:2000 | AEC |
| Monoclonal mouse anti-human BCL6 | Tfh | DAKO, M7211 | 1:50 | AEC |
| **Secondary antibody** | | | | |
| Biotinylated goat anti-rabbit | | DAKO, E0432 | 1:100 | |
| Biotinylated goat anti-mouse | | DAKO, E0433 | 1:100 | |

**Supplemental Table 2: List of the antibodies with their conjugated dye used for Flow cytometric analysis and their working conditions.** All antibodies were purchased from Miltenyi Biotec (except FOXP3, Biolegend). Intercellular targeted antibodies are marked with blue color.

| Channel | PANEL 1 | | Channel | PANEL 2 | |
|---|---|---|---|---|---|
| **V1** | CD3-VioBlue, Cat. 130-114-710 | 1:50 | **V1** | CD3-VioBlue, Cat. 130-114-519 | 1:50 |
| **V2** | Viobility™ 405/520 Fixable Dye, Cat. 130-109-814 | 1µl | **V2** | Viobility™ 405/520 Fixable Dye, Cat. 130-109-814 | 1µl |
| **B1** | CD45-VioBright 515, Cat. 130-110-640 | 1:50 | **B1** | CD25-VioBright 515, Cat. 130-113-287 | 1:50 |
| **B2** | CD20cy-PE, Cat. 130-108-313 | 1:11 | **B2** | BCL6-PE, Cat. 130-118-346 | 1:50 |
| **B3** | CD4-PerCP-Vio700, Cat. 130-113-790 | 1:50 | **B3** | CD4-PerCP-Vio700, Cat. 130-113-228 | 1:50 |
| **B4** | CD8-PE-Vio770, Cat. 130-110-818 | 1:50 | **B4** | CD185 (CXCR5)-PE-Vio770, Cat. 130-117-508 | 1:50 |
| **R1** | Alexa Fluor® 647 anti-human FOXP3 Antibody, Cat: 320114, Biolegend | 1:50 | **R1** | Alexa Fluor® 647 anti-human FOXP3 Antibody, Cat: 320114, Biolegend | 1:50 |
| **R2** | CD68-APC-Vio770, Cat. 130-114-463 | 1:50 | **R2** | CD45-APC-Vio770, Cat. 130-110-635 | 1:50 |

**Supplemental Table 3: Number of cells in the scRNA-seq data sets analyzed in the current study.**

<table>
<tr><td></td><td></td><td colspan="6">Cell number</td></tr>
<tr><td></td><td></td><td>Estimated cell number (cell ranger 10x)</td><td>Primary aggregation (only Donors)</td><td>Secondary aggregation (all samples)</td><td colspan="3">Cell after filtration and normalization</td></tr>
<tr><td rowspan="2">Donor1</td><td>Technical replicate 1</td><td>4276</td><td rowspan="2">7628</td><td rowspan="10">44012</td><td rowspan="2">1592</td><td rowspan="6">4440</td><td rowspan="10">10153</td></tr>
<tr><td>Technical replicate 2</td><td>3352</td></tr>
<tr><td rowspan="2">Donor2</td><td>Technical replicate 1</td><td>3744</td><td rowspan="2">7627</td><td rowspan="2">2748</td></tr>
<tr><td>Technical replicate 2</td><td>3883</td></tr>
<tr><td rowspan="2">Donor3</td><td>Technical replicate 1</td><td>4402</td><td rowspan="2">10033</td><td rowspan="2">2773</td></tr>
<tr><td>Technical replicate 2</td><td>5631</td></tr>
<tr><td>Tumor12MIX</td><td>-</td><td>5878</td><td>-</td><td>2327</td><td rowspan="4">5713</td></tr>
<tr><td>Tumor13SE</td><td>-</td><td>6397</td><td>-</td><td>2440</td></tr>
<tr><td>Tumor2EC</td><td>-</td><td>347</td><td>-</td><td>208</td></tr>
<tr><td>Tumor4SE</td><td>-</td><td>6102</td><td>-</td><td>738</td></tr>
</table>

**Supplemental Table 4: Statistical analysis of different cells of the multiple comparisons across the different localisation of TGCT samples**. The mean of each column was compared with the mean of every other column. Significance tested by ordinary one-way ANOVA including Tukey's Honest Significant Difference Test [(Not significant (ns)= p-value ≥ 0.05), (Significant, *= p-value 0.01 to 0.05, **= p-value = 0.001 to 0.01, ***= p-value 0.0001 to 0.001, ****= p-value <0.0001)].

| | CD3+ cells | CD4+ cells | CD8+ cells | CD68+ cells | CD20cy+ cells | CD25+FOXP3+ cells | CXCR5+BCL6+ cells |
|---|---|---|---|---|---|---|---|
| Seminoma-"Tumor" vs. Seminoma-"Tumor-Adj" | ** (0.0012) | *** (0.0003) | ns (0.3704) | ns (>0.9999) | ns (0.9974) | * (0.0374) | ns (0.2294) |
| Seminoma-"Tumor" vs. Seminoma-"Tumor-Dis" | **** (<0.0001) | **** (<0.0001) | ns (0.0512) | ns (0.9972) | ns (>0.9999) | ns (0.4000) | ns (0.9782) |
| Seminoma-"Tumor" vs. Seminoma-"Contralateral 1" | **** (<0.0001) | **** (<0.0001) | ns (0.0525) | ns (>0.9999) | ns (0.9859) | * (0.0380) | ns (0.8401) |
| Seminoma-"Tumor" vs. Seminoma-"Contralateral 2" | **** (<0.0001) | **** (<0.0001) | * (0.0173) | ns (>0.9999) | ns (0.9227) | ** (0.0022) | ns (0.2904) |
| Seminoma-"Tumor" vs. Embryonal carcinoma (≥80%) –"Tumor" | Ns (0.9896) | Ns (>0.9999) | ns (>0.9999) | ns (0.0666) | ns (0.9989) | ns (0.5582) | ns (0.7885) |
| Seminoma-"Tumor" vs. Mixed Tumors-"Tumor" | Ns (0.1883) | Ns (0.9767) | ns (0.9469) | ns (0.0636) | ns (0.9254) | ns (0.1002) | ns (0.9639) |
| Embryonal carcinoma (≥80%) –"Tumor" vs. Embryonal carcinoma (≥80%)-"Tumor-Adj" | * (0.0180) | ** (0.0052) | ns (0.2172) | ns (0.2668) | ns (>0.9999) | ns (0.9813) | ns (>0.9999) |
| Embryonal carcinoma (≥80%) –"Tumor" vs. Embryonal carcinoma (≥80%)-"Tumor-Dis" | * (0.0281) | ** (0.0018) | ns (0.7592) | ns (0.5719) | ns (>0.9999) | ns (0.9958) | ns (>0.9999) |
| Embryonal carcinoma (≥80%) –"Tumor" vs. Embryonal carcinoma (≥80%)-"Contralateral 1" | * (0.0104) | ** (0.0017) | ns (0.0713) | ns (0.9769) | ns (>0.9999) | ns (0.9974) | ns (>0.9999) |
| Embryonal carcinoma (≥80%) –"Tumor" vs. Embryonal carcinoma (≥80%)-"Contralateral 2" | * (0.0126) | * (0.0216) | ns (0.9910) | ns (>0.9999) | ns (>0.9999) | ns (0.9985) | ns (>0.9999) |
| Mixed Tumors-"Tumor" vs. Mixed Tumors-"Tumor-Adj" | ns (0.8917) | ns (0.4101) | ns (0.9927) | ns (0.2590) | ns (0.3646) | ns (>0.9999) | ns (0.9991) |
| Mixed Tumors-"Tumor" vs. Mixed Tumors-"Tumor-Dis" | ns (0.4644) | * (0.0296) | ns (0.9780) | ns (0.5611) | ns (0.3468) | ns (>0.9999) | ns (>0.9999) |
| Mixed Tumors-"Tumor" vs. Mixed Tumors-"Contralateral 1" | ns (0.2494) | * (0.0262) | ns (0.8948) | ns (0.9750) | ns (0.3842) | ns (>0.9999) | ns (0.9904) |
| Mixed Tumors-"Tumor" vs. Mixed Tumors-"Contralateral 2" | ns (0.2989) | ns (0.1501) | ns (0.8933) | ns (>0.9999) | ns (0.6255) | ns (0.9973) | ns (0.9975) |
